# Are hyaluronic acid synthases widely encoded in fungi?

**DOI:** 10.64898/2026.02.21.705424

**Authors:** Laura Marina Franco-Herrera, Mariandrea Aranda-Barba, Paul Montaño-Silva, Eréndira Patricia Pérez-Muñoz, Jorge Verdín

## Abstract

Hyaluronic acid (HA) is a biologically versatile polysaccharide synthesized by vertebrates and several microbial pathogens. To date, *Cryptococcus neoformans* CPS1p is the only reported hyaluronic acid synthase (HAS) in fungi, which is functionally related to bacterial HASs. Considering the phylogenetic and biochemical connection between chitin synthases (CHSs), essential for fungal cell wall synthesis, and HASs, it is reasonable to hypothesize the latter might be more common in fungi than expected. In this work, a comprehensive *in silico* survey of putative HASs in the Fungal Tree of Life was carried out. 68 putative HASs, mainly in Basidiomycota, were found, although other AI-inferred HASs were found among Ascomycota. Global fold and arrangement of essential amino acids were shared by all kingdoms HASs; however, fungal HASs showed additional exclusive conserved sequence signatures. Moreover, fungal HASs bore an only 3-helices transmembranal pore and their gating loop, which regulates the entrance of substrates to the catalytic site, was directly connected to an also exclusive intrinsically disordered (IDR) C-terminus. Phylogenetically, fungal HASs were found in a clade different to that of bacterial, animal and viral HASs, and all HASs shared the same ancestor with class VI CHSs. The atypical features of fungal HASs could influence the size and biological role of the HA they synthesize and also highlight potential regulatory differences among HASs at the gating loop configuration level.

**Importance:** Despite the report of CPS1p, the hyaluronic acid synthase (HAS) of *Cryptococcus neoformans*, the diversity, structural features and biochemical assets of fungal HASs remain unknown. Here, 68 putative fungal HASs were identified, mainly among Basidiomycota. Although their fold is similar to that of already characterized HASs, their transmembranal pore, integrated by only 3 helices, and their atypical gating loop configuration, suggest they could be also differently regulated, influencing size and function of HA they synthesize.

## Introduction

Hyaluronic acid (HA), also known as hyaluronan, is a linear hetero-polysaccharide composed by alternated units of (1 → 4)-β linked D-glucuronic acid and (1 → 3)-β linked N-acetyl-D-glucosamine^1,2^. HA has been found in vertebrates, some *Streptococcus* and *Pasteurella multocida* bacteria, virus infected *Chlorella*, and the pathogenic yeast *Cryptococcus neoformans*^3–8^. HA plays a variety of roles in each organism. It’s a fundamental component of vertebrates’ extracellular matrix; it is involved in adhesion, proliferation and cell differentiation, and also in the immune response and angiogenesis^9–12^. In microbial pathogens, it is usually a component of a pericellular capsule with two essential functions: adhesion and evasion of the vertebrate host’s immune system^8,13^. Because of its high water retention, high viscosity and biocompatibility, HA is also a relevant biomedical material^14^.

Hyaluronic Acid Synthases (HASs, E.C. 2.4.1.212), the enzymes that catalyze the synthesis of HA, are either transmembranal or peripheric membranal enzymes^15^. For HA synthesis, the enzyme needs UDP-N-acetyl-D-glucosamine (UDP-GlcNAc), UDP-D-glucuronic-acid (UDP-GlcUA), and Mg^2+^ as a cofactor^15,16^. HASs have been classified into two classes^17^: class I HASs bear a catalytic GT2 domain in the cytosolic side of the membrane, a gating loop that regulates the access of substrates to the catalytic site, 3 interphase helices and 4 (bacterial HASs) to 6 (vertebrates and viral HASs) transmembranal helices that form a pore from which synthesized HA is extruded to the extracellular space^18–20^. Class II HASs only contains one member, *P. multocida* HAS. It differs from class I HASs by having two peripheral membranal domains, each one responsible for the catalysis of a different type of glycosidic bond^21^. Class II *P. multocida* HAS can be expressed as a soluble active protein, unlike Class I HAS that must necessarily be associated with the cell membrane^22^.

Most HASs oligomerize. The three isoforms of mammalian HASs form homomeric and heteromeric complexes, but is not clear if they form dimeric, trimeric, tetrameric or higher order oligomeric structures^23,24^. Besides mammalian HASs, *Streptococcus* HAS also forms oligomers^25^. It is possible that oligomers formation is required for HAS activity; moreover, more than one HAS protomer is needed to form the transmembranal pore through which HA in synthesis is extruded^24^. It has also been suggested that the complex formation is necessary to transport the enzyme to the plasma membrane^25^. The dimeric form supports the “pendulum hypothesis” for the mechanism of bacterial HA synthesis^26^: one protomer would bind to the HA chain, while the other binds the UDP-sugar, subsequently, the reaction can proceed^27^.

To date, the pathogenic yeast *C. neoformans* is the sole fungus reported to synthesize HA^8^. There are neither biochemical nor physiological reports of HAS in other fungi, despite it is known that cell walls of *Dactylium*, *Alternaria*, *Fusarium*, *Penicillium*, *Aspergillus*, *Rhizoctonia*, *Neurospora* and *Mucor* contain glucuronic acid^28^. More recently, the same observations were made for *Phycomyces blakesleeanus* and *Rhizopus delemar*^29^. However, the exact nature of the polyuronide they synthesize is unknown.

HAS, like chitin (CHS) and cellulose synthases (CS), are processive beta-glycosyltransferases (GT2 family) characterized by the sequence signature D, D, QXXRW^16,30–32^. Evolutionarily, HASs recently derived from CHSs, central enzymes for the synthesis and morphogenesis of fungal cell walls^17,33^. In support of the latter, it has been observed that HASs are able to synthesize chitin oligomers at lower UDP-GlcUA concentration^16,34^. Considering chitin is a common component of the fungal cell wall, and the biochemical evolutionary connection between CHSs and HASs, it is reasonable to hypothesize a more widespread presence of HASs in fungi. In this work, bioinformatic evidence of the presence of putative Class I HASs in the Kingdom Fungi is provided. The particular characteristics of fungal HASs and their biological implications are discussed.

## Results

HASs have never been described in fungi beyond *C. neoformans*^8^. Nevertheless, the evolutionary and biochemical proximity of HASs to CHSs^17,33^, as well as the report of the synthesis and incorporation of glucuronic acid (HA monomer) into the cell walls of a number of fungi^28,29^ makes reasonable to hypothesize the synthesis of HA, or related polyuronides, in fungi.

### Defining HASs

In order to perform a reliable search of putative HASs in fungi, it was necessary to construct its molecular definition departing from biochemically characterized HASs (Table S1). Because any GT-2 glycosyltransferase can be potentially mistaken as HAS^17,35^, in addition to identify putative HASs, also the fungal orthologs of enzymes that catalyze the synthesis of HA precursors (HAS B, HAS C and HAS D) were identified so that any putative HAS had to be encoded together with HAS B, HAS C and HAS D genes in order to be considered a well supported putative fungal HAS (Figure 1).

**Figure 1.**
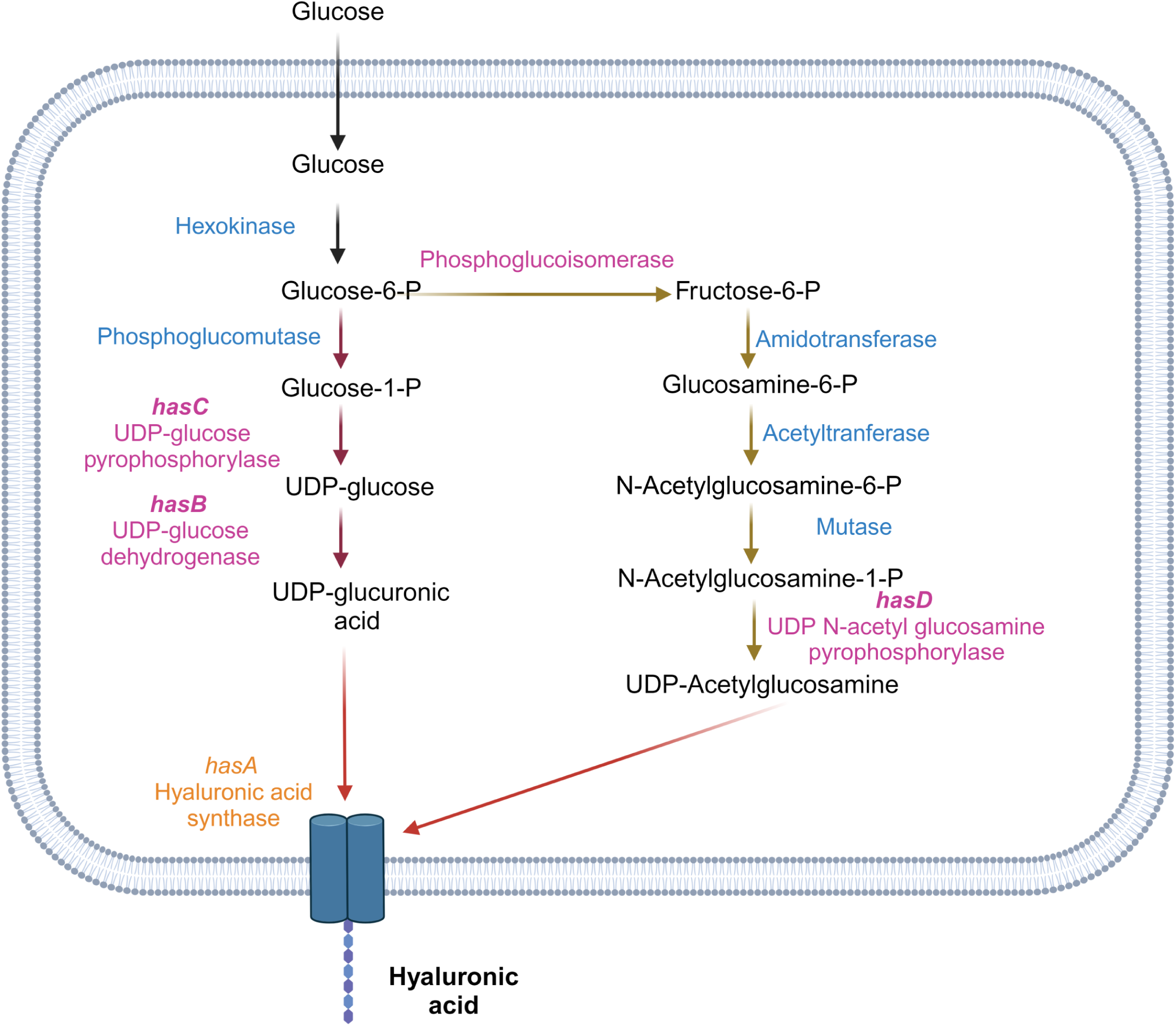
The biosynthetic pathway of hyaluronic acid. The hyaluronic acid biosynthetic pathway derives from glycolysis. From glucose-6-phosphate, the pathway bifurcates to glucose-1-phosphate and fructose-6-phosphate. Glucose-1-phosphate is sequentially transformed to UDP-glucose and UDP-glucuronic acid by UDP-glucose pyrophosphorylase (HAS C) and UDP-glucose dehydrogenase (HAS B), respectively. On the other hand, fructose-6-phosphate is transformed to glucosamine-6-phosphate, N-acetylglucosamine-6-phosphate, N-acetylglucosamine-1-phosphate and, finally, to UDP-N-acetylglucosamine. The latter step is catalyzed by UDP-N-acetylglucosamine pyrophosphorylase (HAS D). Both, UDP-glucuronic acid and UDP-acetylglucosamine are the direct precursors of HA whose synthesis is catalyzed by HAS.

To date, HASs from bacteria (3), vertebrates (23), a virus and a yeast have been identified and fully or at least partially characterized (Table S1A)^4–8,36–38^. In addition, essential residues have been identified for a number of HASs^18,39^ (Figure S1). Sequences of 28 biochemically characterized HASs were retrieved (Table S1A) and bioinformatically analyzed. Sequence size spanned from 417 to 588 residues. 64 amino acids were highly conserved; 30 of these residues have been previously studied and their function in the activity of *Streptococcus equi* HAS (SeHAS) is known^18,39^. The function of the other 34 residues is unknown. However, their importance was inferred from their high conservation along all analyzed sequences (>90%) (Figure S1 and Table S2).

Among conserved residues, some of them are definitory of GT-2 family glycosyltransferases: QXXRW, WGTR and DXD (Table 1). On the other hand, conserved signatures KR, GDD, SG and LT are characteristic of HASs, as well as residues G38, I71, E76, S87, G186, N192, R205, E213, C226, Y233, F250, G252, G271, R305, R368, P396 and T404 (SeHAS numbering, see Table S2 for numbering equivalence in *C. neoformans* CPS1p, CnHAS). Previous reports suggested an important role for those residues in protein stability (SG), substrate recognition (GDD) and processivity (KR)^18^. Additional signatures, exclusive to HASs of specific Kingdoms, were found (Figure S2). Bacteria, vertebrates and viruses HASs conserved KS and RY motifs. Instead of RY, yeast’s CnHAS harbored an RL motif. Bacteria and vertebrates HASs conserved an AF motif. Vertebrates and virus HASs shared a TXL motif, in bacteria TXI. Vertebrates harbored FA, WL and GXVGG signatures, the latter also present in yeast’s CnHAS as a GGVG motif (Figure S2).

**Table 1.** Conserved motifs that define fungal hyaluronic acid synthases.

| GT-2 motifs | General HASs motifs | Exclusive fungal HASs motifs |
| --- | --- | --- |
| QXXRW | KR | GGVXT |
| WGTR | SG | WEXL |
| DXD | GDD | RTAXY |
|  | LT | LXD |
|  | G38, I71, E76, S87,<br>G186, N192, R205, E213,<br>C226, Y233, F250, G252,<br>G271, R305, R368, P396<br>and T404 | FXXRW |
|  |  | WXXXP |

**Table 2.** Sub-classification of class I hyaluronic acid synthases.

| Subclass | α-HAS | β-HAS | γ-HAS |
| --- | --- | --- | --- |
| **Kingdom** | Vertebrates and viral HASs. | Bacterial HASs. | Fungal HASs. |
| **Gating loop configuration** | Connected to IFH3 and TMH5 (transmembranal pore). | Connected to IFH3 and finishes at C-terminal end. | Connected to IFH3 and C-terminal IDR. |
| **Transmembranal pore helices** | 6 | 4 | 3 |
| **Oligomerization state** | Vertebrates HASs can be monomeric or oligomeric. Viral HAS is monomeric. | Oligomeric. | Not determined yet. |
| **HA elongation** | Non-reducing end | Reducing end | Not determined yet. |

### Putative fungal HASs are more abundant, but not exclusive, to Basidiomycota

Departing from sequences of already characterized HASs, HMM’s were constructed and used to insightfully search for putative HASs in the inferred proteomes of 910 fungal species representatives of the Fungal Tree of Life. This approach allowed the search of putative HASs with a wide range of sequence variation, but restricted to maintain essential motifs for HAS activity (Table 1 and S2, Figure S1). According to this strategy, 68 putative fungal HASs were retrieved from the analyzed fungal proteomes database, belonging to 64 species (Table S3). Most of them encoded a single HAS, but S*ynchytrium microbalum*, *Lobosporangium transversale*, *Phycomyces blakesleeanus* and *Suillus fuscotomentus* encoded two HASs. Putative HASs were found in Ascomycota (2), Basidiomycota (57), Chytridiomycota (4) and Mucoromycota (5) (Figure 2). The only two HASs of Ascomycota belonged to Eurotiales species. Among Basidiomycota, putative HASs were found mainly in Tremellales, Boletales, Agaricales and Polyporales species (Figure 2), while in Mucoromycota they mostly appeared in Mucorales and Mortierellales species. Finally, among Chytridiomycota, putative HASs were found encoded in Rhizophydiales, Spizellomycetales and Synchytriales species (Figure 2).

**Figure 2.**
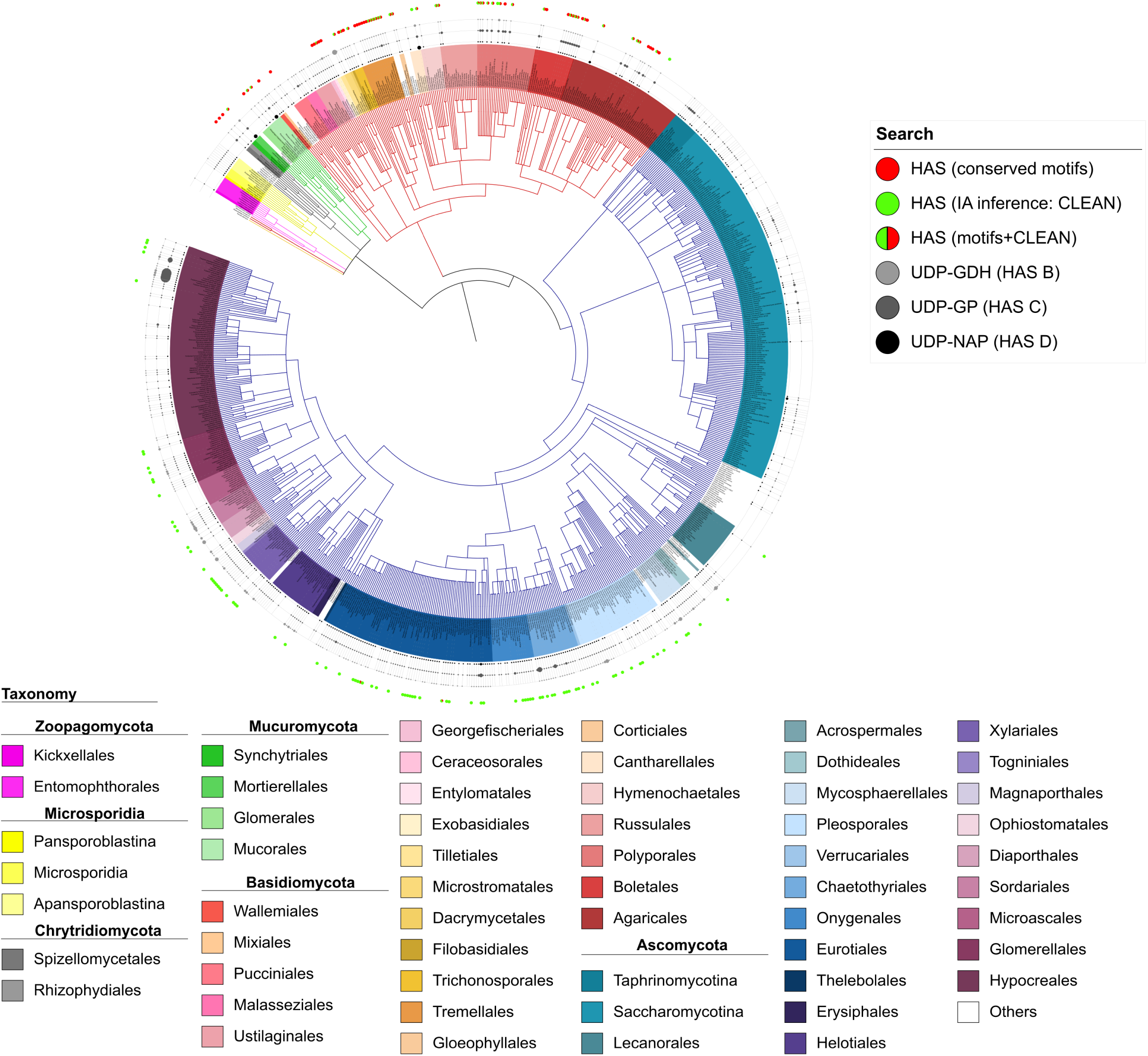
Putative hyaluronic acid synthases in the Fungal Tree of Life. Putative fungal orthologs of HAS were retrieved from a 910 fungal species proteomes database using HMMs based on already biochemically characterized HASs plus conservation of typical HASs motifs, or inferred with the artificial intelligence tool, CLEAN^40^. To further support fungal HASs prediction, species encoding a putative HAS were assessed to also encode HAS B (UDP-glucose-6-dehydrogenase), HAS C (UDP-glucose pyrophosphorylase) and HAS D (UDP-N-acetylglucosamine pyrophosphorylase), all them enzymes of the HA synthetic pathway. HMMs+conserved motifs-inferred fungal HASs mostly appeared in Chytridiomycota, Mucoromycota and Basidiomycota, and the species that encoded them consistently also encoded HAS B, C and D. Results of putative fungal HASs prediction were grafted into a fungal tree of life constructed on the 910 reference fungi database using PhyloT and based on the NCBI taxonomy^80^. The final figure was created with iTOL^81^.

To further support the search, putative fungal HASs were also inferred with CLEAN, an artificial intelligence tool that uses contrastive learning to assign a function to uncharacterized enzymes, with a better accuracy than traditional tools^40^. CLEAN results mostly coincided with that of HMM + conservation of typical HASs motifs-based search (Table S3); however, it also identified abundant putative HASs among Ascomycota. Within this division, HASs were mostly predicted in Pleosporales, Chaetothyriales, Onygenales, Eurotiales, Xylariales and Glomerellales species (Figure 2). Despite CLEAN unambiguously assigned EC 2.4.1.212 (hyaluronic acid synthase) activity to these enzymes, the F1 factor that assessed the reliability of the prediction was consistently low (F1<0.6).

As stated above, it was realized that any *bona fide* HAS should be escorted by the enzymatic machinery necessary to synthesize HA precursors; thus, a parallel search in the 910 fungal proteomes database was performed for HAS B, HAS C and HAS D orthologs. These enzymes are part of the HA biosynthetic pathway shared by bacteria, fungi and animals (Figure 1)^41,42^. HAS B (UDP-glucose dehydrogenase) catalyzes the transformation of UDP-glucose into UDP-glucuronic acid, HAS C (UDP-glucose pyrophosphorylase) synthetizes UDP-glucose from glucose-1-P, and HAS D (UDP N-acetylglucosamine pyrophosphorylase) produces UDP-GlcNAc from glucosamine-1-P and acetyl-CoA^41,42^.

Already biochemically characterized HAS B, C and D (Table S1B-D) were used to build HMMs to retrieve their fungal orthologs. Orthology was parallelly confirmed by CLEAN and phylogenetic analysis. In the case of HAS B, 4 over 5 reference HAS B (XP_012048628.1, ABW95825.1, XP_006666904.1 and XP_040878575.1) got a confirmatory CLEAN prediction (E.C. 1.1.1.22, UDP-glucose-6-dehydrogenase) and the inferred orthologs within their clades in HAS B phylogenetic reconstruction were also considered HAS B (Figure S3). Nevertheless, despite *A. fumigatus* XP_753070.1^43^, the 5th reference HAS B, was also functionally confirmed, only obtained a CLEAN F1=0.46, well below the stipulated 0.6 cutoff. Because the activity of *A. fumigatus* XP_753070.1 had been empirically demonstrated^43^, the CLEAN prediction was taken as correct and the sequences of the entire clade where XP_753070.1 clustered were considered also HAS B. After the screening, 347 HAS B orthologs were identified in 300 species from all fungal phyla (Figure S3).

For HAS C orthologs, reference HAS C used to retrieve them were functionally confirmed by CLEAN (E.C. 2.7.7.9, UDP-glucose pyrophosphorylase; F1>0.6), which extended to all retrieved HAS C orthologs. 534 fungal HAS C orthologs were identified in 454 species. They were widely allocated in the fungal tree of life (Figure S4).

In the phylogeny reconstruction of the first set of HAS D orthologs retrieved with a HMM based on five reference HAS D, two main clades were observed (not shown). For one of the clades, CLEAN predicted the expected E.C. 2.7.7.23 (UDP-N-acetylglucosamine pyrophosphorylase), but also E.C. 2.7.7.83 (UDP-N-acetylgalactosamine diphosphorylase). It has been reported that animals HAS D can harbor both E.C 2.7.7.23 and E.C. 2.7.7.83 activities^44,45^; thus, the sequences of this clade were preliminary considered HAS D orthologs. In the other clade were identified mostly L-rhamnonate dehydratases (E.C. 4.2.1.90), followed by gluconate dehydratases (E.C. 4.2.1.39), ferredoxin oxidoreductases (E.C. 1.3.7.5), cysteine synthases (E.C. 2.5.1.47), alanine-glyoxylate transaminases (E.C. 2.6.1.44), and homospermidine synthases (E.C. 2.6.1.45), which were all discarded. The sequences of the first clade were required to also contain the nucleotide binding motif LXXGGQGTRLGXXXPK that defines UDP-N-acetylglucosamine pyrophosphorylases^46,47^. As a result, 99% of the parsed sequences were functionally predicted as E.C. 2.7.7.83 by CLEAN. The phylogeny of the remaining 449 sequences was reconstructed and shown in Figure S5. HAS D appeared in 440 species within the main fungal phyla.

All the species that encoded HMM+conservation of typical HASs motifs-inferred HASs, also encoded the three enzymes involved in HA synthesis: HAS B, C and D, within Chytridiomycota, Mucoromycota, Basidiomycota and Ascomycota (Figure 2). Even CLEAN predicted HASs that mostly populated Ascomycota species, were escorted by HAS B, HAS C and HAS D encoding genes. However, these putative HASs did not show at least one of the motifs critical for catalytic activity; that is why this set of putative HASs was discarded in the following analysis. Ascomycetes that did not encode HMM nor CLEAN predicted HASs genes were also devoid of HAS B, although they usually encoded HAS C or HAS D.

Besides the generally conserved motifs among HASs (Table 1, Figure S2), putative fungal HASs also showed motifs exclusively conserved among them: V[T/S]53, PTI58, [W/M]L73, P78, T85, N114, Q[M/L]118, G122, W140, L149, GGVXT158, WEXL176, RN187, G199, L204, RTAXY207, LXD216, F[T/Q]225, FXXRW243, W253, Q258, S289, WXXXP301, H370 and F391 (CnHAS numbering) (Figure S2). These motifs have not been studied in CnHAS; however, due to its high conservation, they could play essential roles in the stability or catalysis of this enzyme.

### The catalytic domain of different Kingdoms HASs is structurally conserved

To further support the inference of new putative fungal HASs, the predicted structure of CnHAS catalytic domain, to date the only *bona fide* fungal HAS^8^, and a good representative of the group, was structurally aligned to those of *S. equi subsp. zooepidemicus* HAS (SeHAS), *Homo sapiens* HAS2 (HsHAS2) and *Xenopus laevis* HAS (XlHAS), well representatives of bacteria and animal HASs. The three different Kingdoms HASs catalytic domains aligned with an RMS of 1.047 Å (CnHAS *vs* SeHAS), 1.137 Å (CnHAS *vs* HsHAS2), 1.141 Å (CnHAS *vs* XlHAS), 1.106 Å (SeHAS *vs* XlHAS), 0.95 Å (HsHAS2 *vs* XlHAS) and 0.835 Å (SeHAS *vs* HsHAS2), confirming all different Kingdoms HASs catalytic domains shared the same general fold (Figure 3A). This fold conservation was also observed among the catalytic domains of representative putative fungal HASs, *Coprinopsis cinerea* HAS (CcHAS) and *Mucor circinelloides* HAS (McHAS), in comparison to CnHAS (CnHAS *vs* CcHAS, 0.505 Å RMSD; CnHAS *vs* McHAS, 0.583 Å RMSD; and CcHAS *vs* McHAS, 0.552 Å RMSD) (Figure 3B).

**Figure 3.**
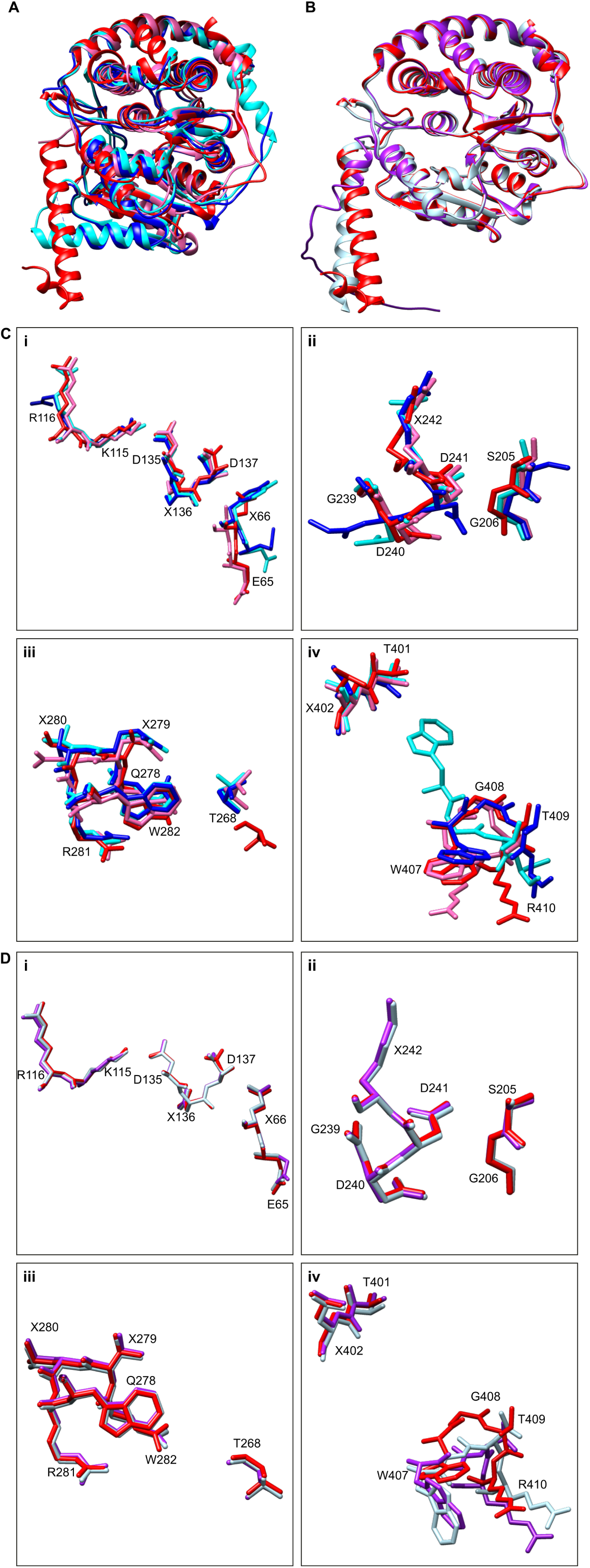
Structural conservation of different Kingdoms HASs catalytic domains. A) CnHAS (red) catalytic domain shared the same fold as SeHAS (pink), XlHAS (blue) and HsHAS2 (cyan). B) The fold of CnHAS catalytic domain (red) was fully conserved among representative putative fungal HASs, CcHAS (light blue) and McHAS (purple). C) Structural alignment of the catalytic motifs of CnHAS, SeHAS, XlHAS and HsHAS2, and D) CnHAS, CcHAS and McHAS (same colors as described above). C and D, i) structural alignment of the motifs KR and EX important for HAS activity and DXD characteristic of glycosyltransferases GT2; ii) structural alignment of the motifs GDDX and SG, also involved in the activity of HAS; iii) alignment of the motif QXXRW shared by all the processive glycosyltransferases of the family GT2, and residue T268, relevant for HASs catalysis; iv) alignment of motifs TX and WGTR; the latter is characteristic of the gating loop. The models for CnHAS, SeHAS, HsHAS2, CcHAS and McHAS were built with AlphaFold-3^89^. The structure of XlHAS was obtained from the Protein Data Bank (PDB ID: 8smn). The alignment and edition of structures were made in Chimera 1.18^95^.

Likewise, the previously described catalytic motifs, KR, DXD, EX (Figure 3Ci, Di), GDDX, SG (Figure 3Cii, 3Dii), QXXRW, T268 (Figure 3Ciii, Diii), TX and WGTR (Figure 3Civ, Div) of the biochemically characterized HASs (CnHAS, SeHAS, HsHAS2, and XlHAS) were structurally aligned (Figure 3C), as well as those of CnHAS with homologous motifs of the putative fungal HASs of *M. circinelloides and C. cinerea* (Figure 3D). Besides the motifs conservation, their geometrical arrangement is also conserved in all Kingdoms HASs.

Class I HASs bind both substrates in a single catalytic site^18^. Molecular docking analysis of AlphaFold modeled CnHAS with HA precursors, UDP-GlcNAc (−9.9 kcal/mol) and UDP-GlcUA (−9.8 kcal/mol) (Figure 4A-C), confirmed both substrates interact with almost the same amino acids in CnHAS catalytic site, homologous to those reported for other Kingdoms HASs (Figure 4 and S6)^18^. GlcNAc interacted with L184, R187, S205, R207, D241, K242, W282 (Figure 4D), while GlcUA also interacted with the previously mentioned residues (except K242) and Q164 (Figure 4E). The UDP moiety of the substrate sugars interacted with the motif DDD (residues 135-137), M269, R281, W407 and R410 (Figure 4D and E). The latter two amino acids are part of the gating loop (Figure 4A, D, E). A negative control docking analysis with R281A/W282A CnHAS mutant, which replaced RW amino acids in the substrate binding motif QxxRWxKSxxRE, and both substrates, was also performed (Figure S6). As expected, the affinity energies depleted for both UDP-GlcNAc (−8.1 kcal/mol) and UDP-GlcUA (−7.9 kcal/mol). In addition, the amino acids that interacted with the substrates changed: with UDP-GlcNAc interacted D61, E65, K115, 135DDD137, R207, T268, K270, Q278, W407 and R410; with UDP-GlcUA interacted almost the same amino acids as UDP-GlcNAc, except E65, K115, R207 and T268; additionally, this substrate interacted with M269. These results indicated the validity of the above-described docking analysis with wild type CnHAS. The type of interaction between the UDP-sugars and each amino acid of wild and mutant CnHAS catalytic pocket can be seen in Table S4.

**Figure 4.**
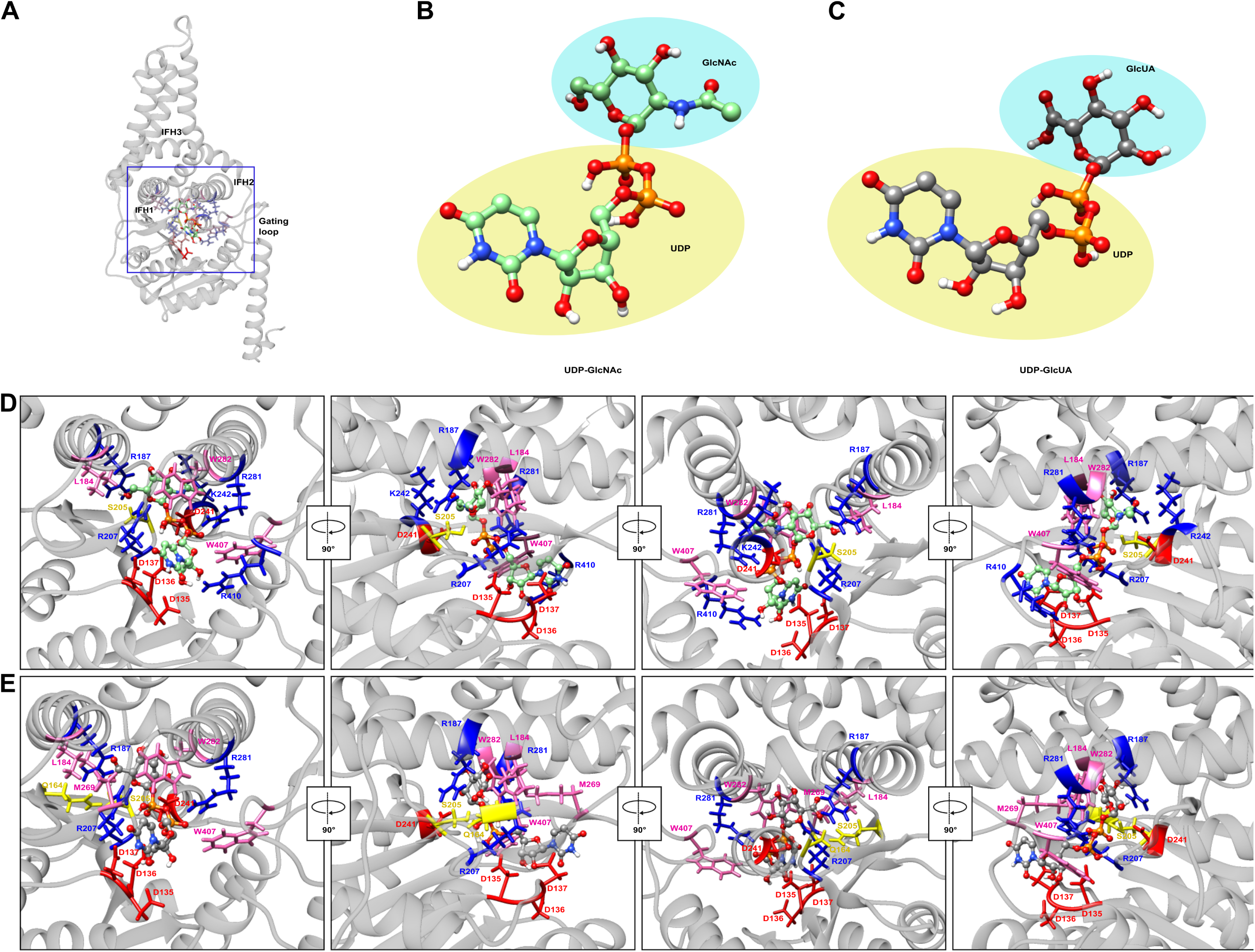
Molecular docking analysis of Cryptococcus neoformans HAS and its substrates, UDP-GlcNAc and UDP-GlcUA. (A) *C. neoformans* HAS structure was modeled with AlphaFold 3^89^ and used to perform molecular docking analysis in Chimera 1.18 using Autodock Vina^95^ to determine the interaction features with (B) UDP-GlcNAc and (C) UDP-GlcUA. D) Orthogonal views (90 ° rotation on “Y” axis) of the molecular docking analysis between modeled *C. neoformans* HAS and UDP-GlcNAc as substrate. UDP-GlcNAc interacted with 135DDD137, L184, R187, S205, R207, D241, K242, R281, W282, W407 and R410. E) Orthogonal views (90 ° rotation on “Y” axis) of molecular docking analysis between *C. neoformans* HAS and UDP-GlcUA as substrate. UDP-GlcUA interacted with 135DDD137, Q164, L184, R187, S205, R207, D241, M269, R281, W282 and W407.

### The transmembranal pore of fungal HASs was formed by only three alpha-helices

Despite modeled 3D structures of *C. neoformans*, *S. equi*, *H. sapiens*, and *X. laevis* HASs were structurally homogeneous conserving the typical GT-2 catalytic domain (DXD, QxxRW conserved motif in a ‘finger helix’ dynamically coupled with the WGTR gating loop motif)^35^ and the transmembranal pore from which the growing polysaccharide is extruded to the extracellular milieu^18,48^, the composition of the latter was conspicuously variable (Figure 5A). The transmembranal pore of bacterial HAS (SeHAS) was formed by 4 helices (H1, H2, H3 and H4; Figure 5A), while the pore of animal and viral (*H. sapiens*, *X. laevis* and CvHAS) HASs was composed of 6 helices (H1, H2, H3, H4, H5 and H6; Figure 5A). In contrast, the transmembranal pore of fungal HASs (CnHAS) was formed exclusively by 3 helices (H2, H3 and H4; Figure 5A). Thus, bacterial HASs were devoid of helices H5 and H6, while fungal HASs lacked helices H1, H5 and H6 (Figure 5A). Besides transmembranal helices, the HAS pore was also formed by three interface helices (IFH1, IFH2, IFH3) that were strictly conserved in all HASs analyzed (Figure 5B). The IFH helices of CnHAS preserved the structure, spatial location and hydrophobicity pattern of those of SeHAS, XlHAS, HsHAS2 and CvHAS (Figure S7).

**Figure 5.**
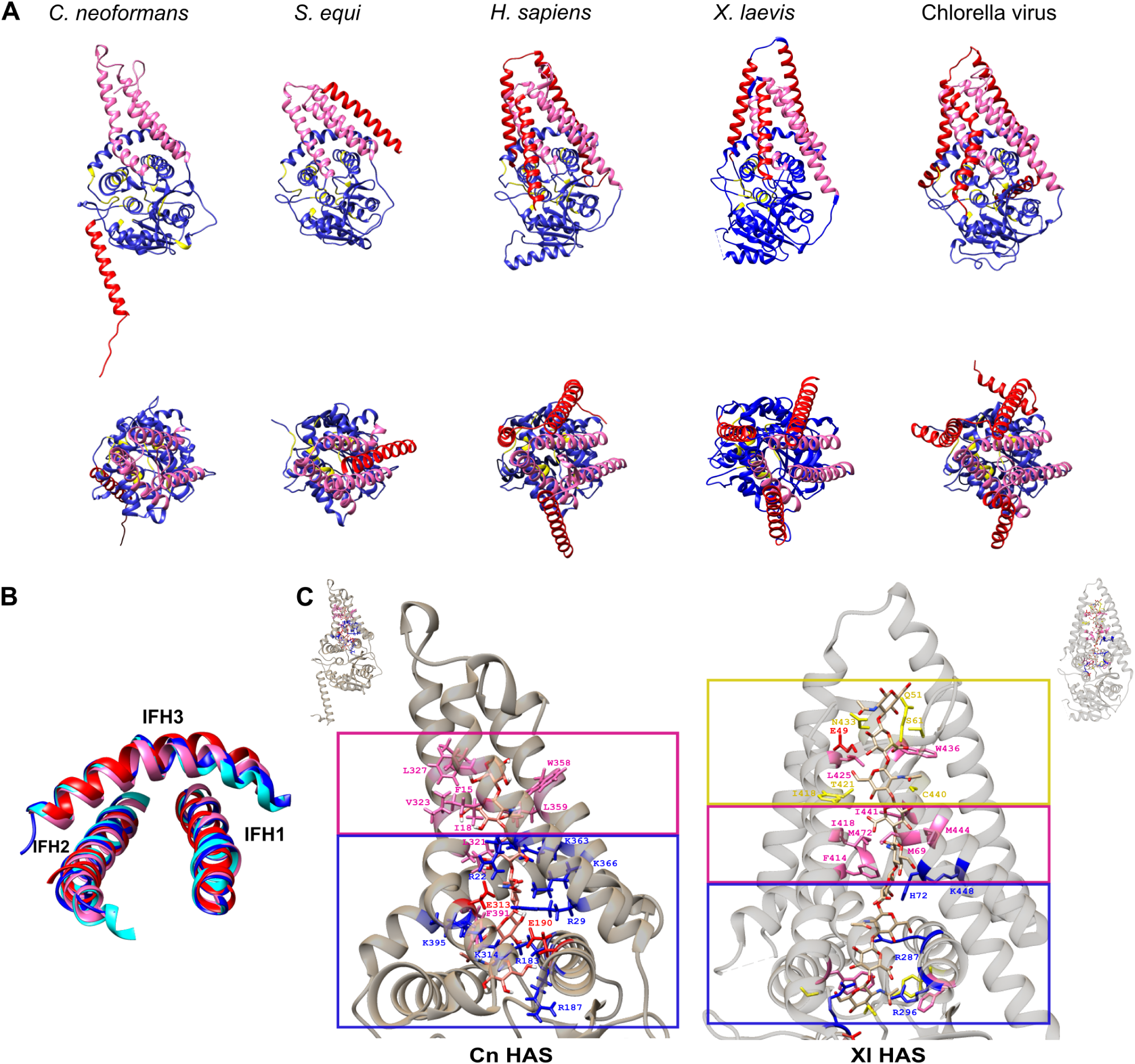
The structural diversity of HAS transmembranal pore. A) Fungal (CnHAS), bacterial (SeHAS), animal (HsHAS2, XlHAS) and viral (CvHAS) HASs share the same general fold. However, the pore of CnHAS, and all putative fungal HASs, is formed by only 3 transmembranal helices, which contrasted with the pore of bacterial HASs (SeHAS) formed by 4 transmembranal helices, and vertebrates (HsHAS2, XlHAS) and viral HASs (CvHAS) formed by 6 transmembranal helices. HASs structures were modeled in AlphaFold 3^89^ and displayed with Chimera 1.18^95^. Upper row, lateral view of HASs structures where the catalytic domain is blue colored, while the helices of the pore are colored in pink (shared by fungal HASs) and red (absent in fungal HASs). Lower row, cenital view of HASs transmembranal pore. B) CnHAS (red) conserved the 3 interface helices (IFH1-IFH3) which structurally align with the IFH of SeHAS (pink), XlHAS (blue) and HsHAS2 (cyan). C) The docking analysis on CnHAS with a HA hexasaccharide showed that the pore (left), formed by 3 transmembranal helices and 3 interface helices, had the same biochemical characteristics of that of XlHAS (ID: 8smn, right) that facilitates the translocation of HA. In the base of the pore a ring rich in positively charged amino acids (blue square, R22, R29, R183, R187, R314, R363, K366 and K395) was found, followed by a collar of hydrophobic amino acids (pink square, F12, F15, I18, L321, V323, L327, W358). These features are consistent with those of the XlHAS pore (right); however, CnHAS pore lacked the last ring described for XlHAS^48^, which contains hydrophobic and polar amino acids (yellow square). Docking simulations and subsequent analyses were performed in Chimera 1.18 using Autodock Vina^95^, with an HA hexasaccharide obtained from GLYCAM (<u>glycam.org</u>).

To assess any change in the properties of fungal HASs transmembranal pore due to their different structural configuration, a molecular docking analysis on CnHAS was performed (Figure 5C). A hexasaccharide of HA managed to couple (−10.5 kcal/mol) in the pore of CnHAS (Figure 5C, left), despite the lack of H1, H5 and H6 helices. In the base of the pore, the HA oligomer interacted with a collar rich in positively charged amino acids (R22, R29, R183, R187, K314, K363, K366, K395), followed by a ring of hydrophobic amino acids (F15, I18, L321, L327, V323, W358, L359). In a docking analysis between the R29A/D313A mutant CnHAS and HA oligomer (−9.6 kcal/mol), used as negative control, HA did not manage to couple the pore (Figure S8). Compared with the interaction model of XlHAS pore with an HA oligomer (Figure 5C, right; PDB ID: 8smn)^48^, CnHAS showed only 2 out of 3 rings of those identified in XlHAS pore. CnHAS lacked the third ring at the end of the pore close to the extracellular milieu, which is rich in polar and hydrophobic amino acids. Even though the 2 collars that CnHAS conserved did not retain the amino acids sequence identity, they conserved the biochemical characteristics.

### The gating loop is strictly conserved among different Kingdoms HASs, but it is differently connected to other elements of the protein

The gating loop is a structure conserved in all HASs reported so far. This cytosolic loop interacts with the nucleotide bound to the substrate sugar, opening the way to the catalytic site and triggering the synthesis and translocation of HA^48^. The CnHAS gating loop had a similar structural configuration as the bacterial gating loop (SeHAS, Figure 6A). However, although both CnHAS and bacterial gating loops came after the IFH3, bacterial HASs gating loop was at the very C-terminus of the protein, while CnHAS gating loop was localized at the beginning of a C-terminal intrinsically disordered region (IDR) (Figure 6A). Disorder prediction showed the C-terminal IDR was exclusive and conserved along all the putative fungal HASs (Figure 7), with exception of Ascomycota HASs that, in addition, harbored a slightly divergent gating loop motif, WGSR. The size of these IDRs was between 30 and 200 amino acids, depending on the protein size (Figure 7). In contrast, vertebrate’s HASs (Figure 6B-C) gating loops were C-terminally connected to an alpha helix (TMH5) that formed part of the transmembranal pore, adopting a U-like configuration. In addition, animal HASs gating loop motif was longer, but clearly conserved (WGTSGRK/R, Figure S9) between IFH3 and TMH5 helices^48^. CvHAS conserved the WGTR gating loop motif, mentioned before for CnHAS and bacterial HASs (Figure S9). Despite divergence of gating loop motifs and differential structural connection, the gating loop was conformationally conserved in all Kingdoms HASs (Figure 6).

**Figure 6.**
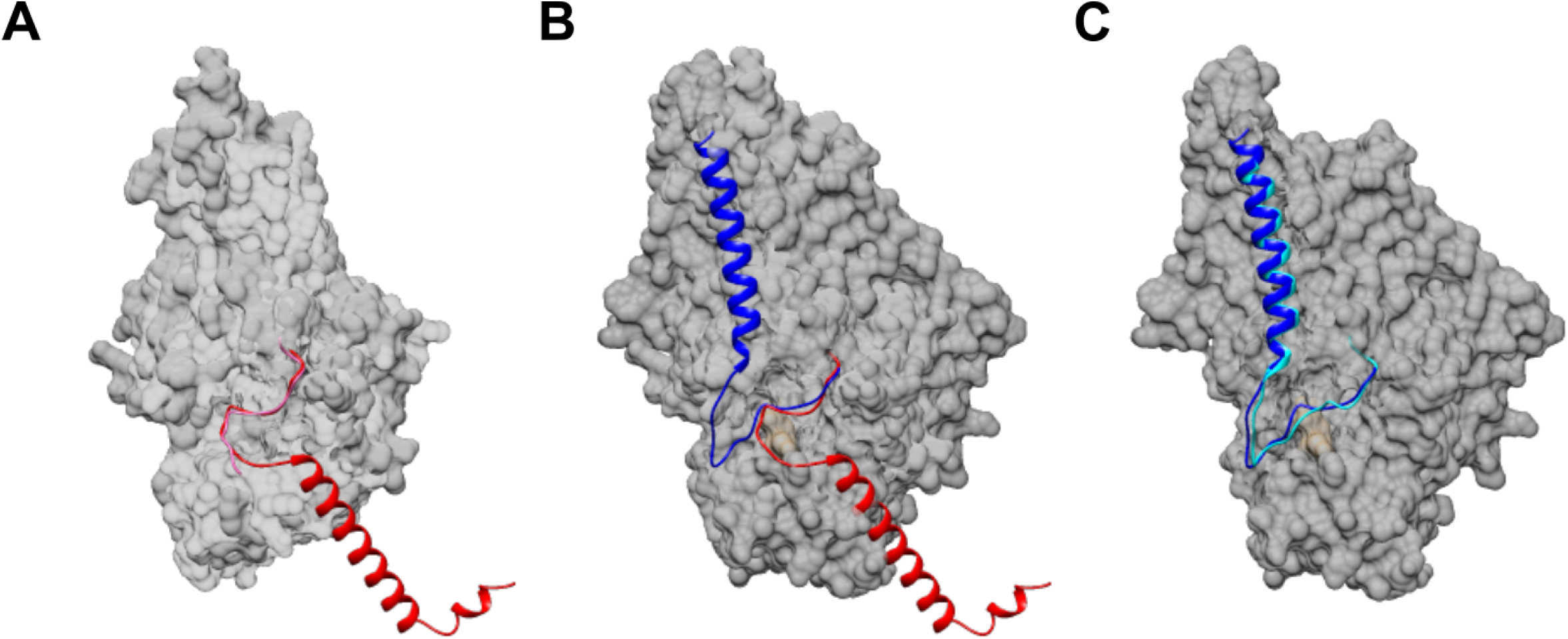
The gating loop is strictly conserved among different Kingdoms HASs, but it is differently connected to other elements of the protein. A) The gating loop of CnHAS (red) structurally aligns with the gating loop of SeHAS (pink). Nevertheless, CnHAS gating loop is connected to an intrinsically disordered region (IDR). B) In contrast, XlHAS gating loop (blue) is connected to H5, which forms part of the transmembranal pore. Similarly, C) HsHAS2 gating loop (cyan) is connected to a helix that also forms part of the pore. 3D structures of CnHAS and HsHAS2 were modeled with AlphaFold 3^89^, while XlHAS structure was taken from the Protein Data Bank (PDB ID: 8smn). Structural alignment and surface display were performed in Chimera 1.18^95^.

**Figure 7.**
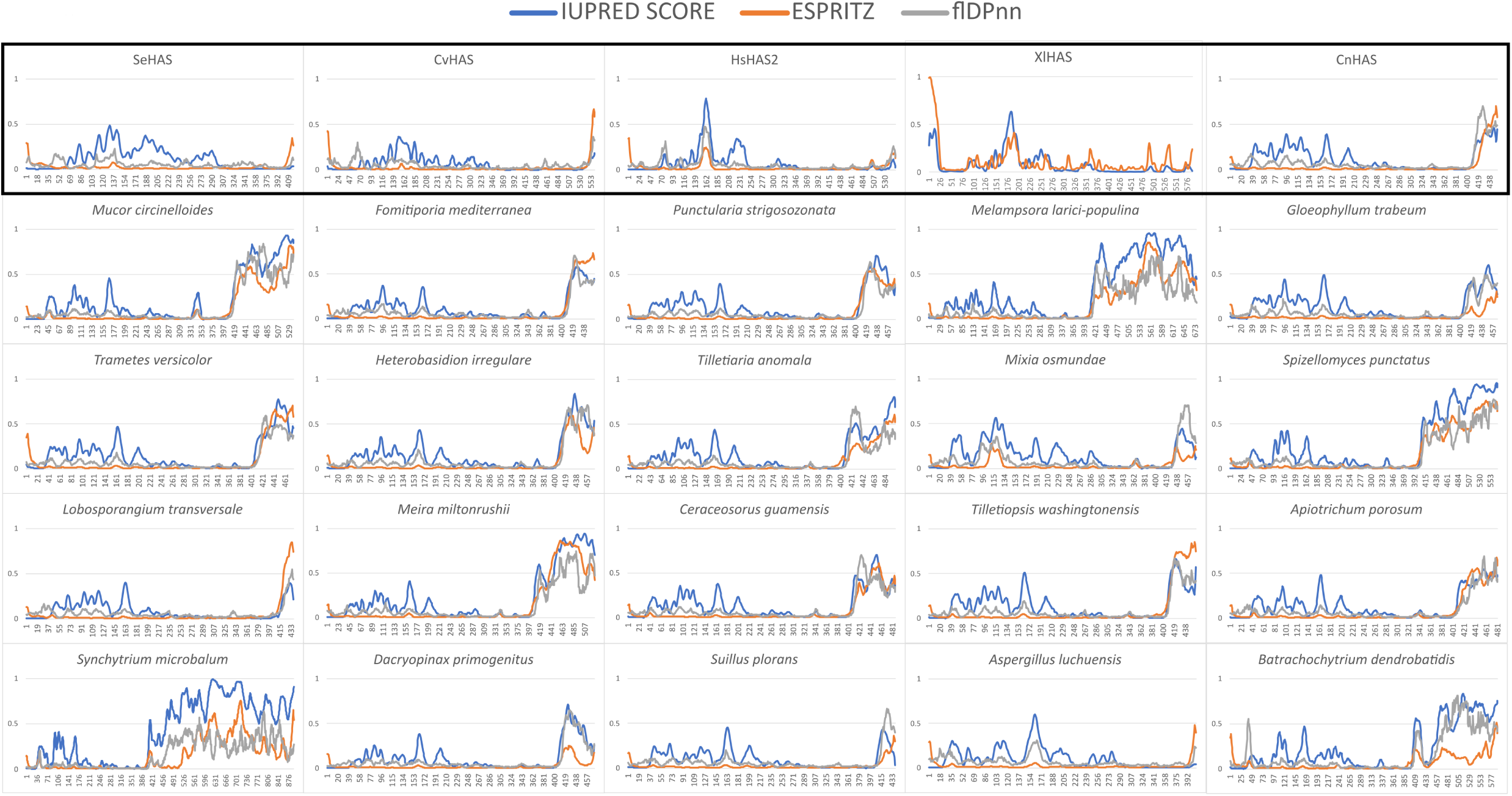
Fungal HASs are intrinsically disordered at the C-terminus. In contrast to bacterial (SeHAS), viral (CvHAS) and animal HASs (HsHAS2 and XlHAS), CnHAS C-terminus, just after the gating loop, harboured an intrinsically disordered region (IDR). This feature was consistently predicted in a representative HAS of each fungal order that contained one. The average size of this intrinsically disordered region was between 30 and 200 amino acids, which suggested it could be biologically relevant. Intrinsic disorder was predicted in three platforms: IUPRED3^91^, ESpritz 1.3^92^ and flDPnn^93^.

### Putative fungal HASs clustered homogeneously together, but differentiated from bacterial, viral and vertebrates HASs

The phylogenetic reconstruction of the 68 putative fungal HASs showed a distribution that did not parallel the speciation events within Kingdom Fungi, which indicates an independent evolution of putative fungal HASs (Figure 8A). Ascomycota, Chytridiomycota and Mucoromycota putative HASs appeared in the same clade (Group I). On the other hand, Basidiomycota HASs segregated in two different clades: Group II, which encompassed HASs from Georgefisheriales, Ceraceosorales, Entylomatales, Exobasidiales, Trichosporonales and Tremellales, and Group III, which included putative HASs from Dacrymycetales, Gloeophyllales, Polyporales, Boletales, Hymenochaetales, Agaricales, Russulales, and Corticiales (Figure 8A).

**Figure 8.**
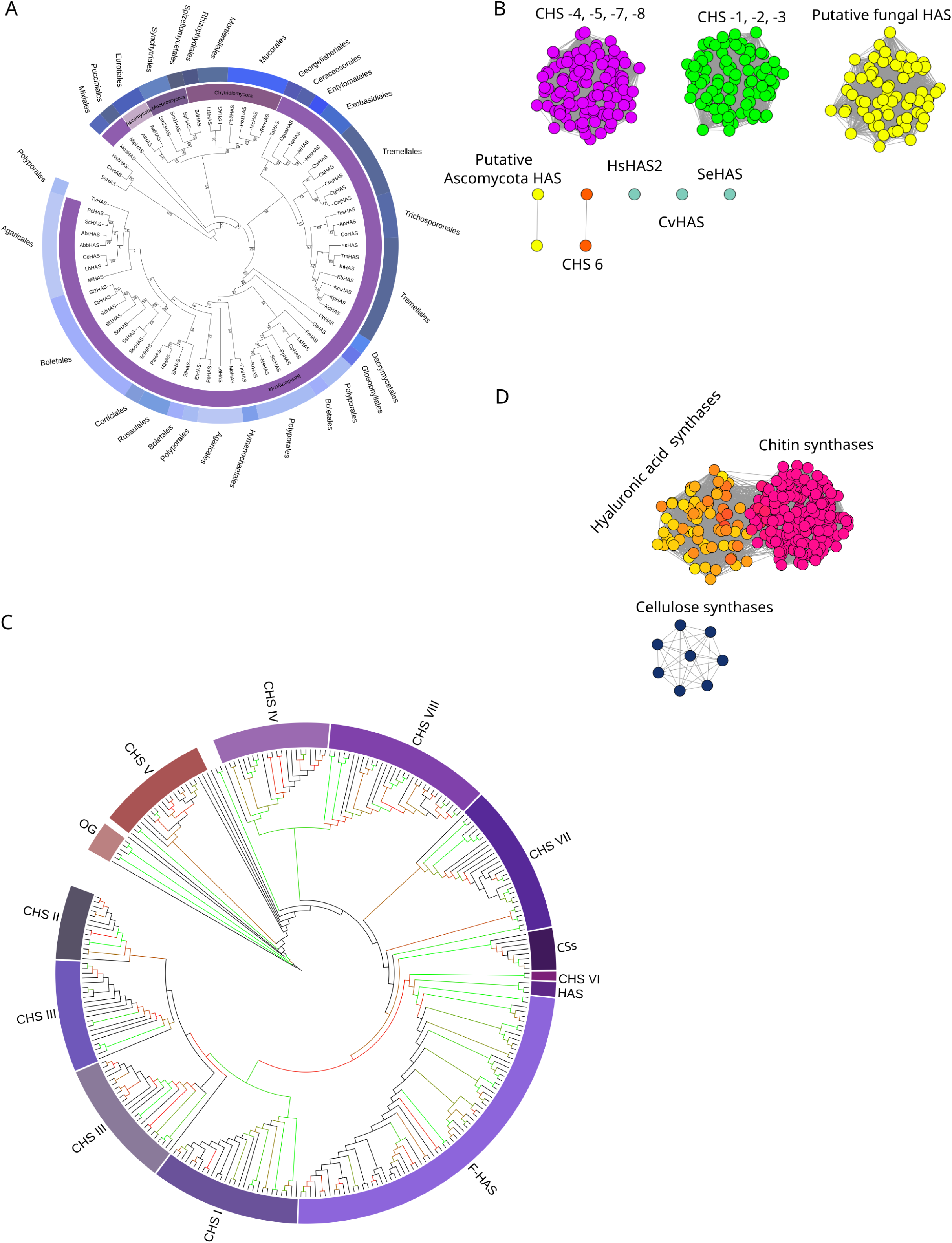
*Phylogenetic reconstruction of putative fungal HASs* and their relationship with other Kingdoms HASs, chitin and cellulose synthases. A) Phylogenetic reconstruction of putative fungal HASs, where they were clustered in three main clades: Chytridiomycota, Mucoromycota and Ascomycota HASs are arranged together in group I, while Basidiomycota HASs sorted out in group II and III. Bootstrap values are indicated at the node. SeHAS, HsHAS, and CvHAS were used as outgroups. B) Sequence similarity network (SSN) analysis using the 68 fungal HASs and representative chitin synthases (CHSs) sequences from the eight classes. CHSs were clustered in three different groups: class IV, V, VII and VIII CHSs were clustered together, while class I, II and III CHSs were put in the same group. Only class VI CHSs, fungal HASs and putative fungal HASs from Ascomycota were segregated in their own cluster. Reference HsHAS2, SeHAS, and CvHAS behaved as singletons. The same arrangement was observed in the phylogenetic reconstruction (C), where cellulose synthases (CS) from plants (*A. thaliana*) and bacteria were also included. Bacterial CSs were used as an outgroup. Bootstrap values from 50 to 100 are displayed in color gradient from minimum (50) to green (100). D) SSN analysis using CHSs, fungal HASs and CSs, support the idea that CHSs are more closely related to fungal HASs than CSs.

Already characterized HASs from bacteria (SeHAS), animals (HsHAS2) and virus (CvHAS) located in a different clade than fungal HASs (Figure 8A). Clustering analysis of reference and putative fungal HASs confirmed the segregation of the latter into a compact nucleus (Figure 8B). Nevertheless, the two Ascomycota HASs clustered apart from the main cluster, probably because of the lack of the C-terminal IDR and changes in important motifs sequence, such as the gating loop motif (WGSR instead of the more prevalent WGTR). As expected, bacterial (SeHAS), viral (CvHAS) and vertebrate (HsHAS2) HASs neither clustered with the fungal ones nor between them, which outlined their differences despite all them harbored the signatures that make them *bona fide* HASs (Figure 8B).

### Fungal HASs are evolutionary closer to class VI chitin synthases

Some authors have suggested the evolutionary connection between HASs and CHSs^17,33,49,50^. The capacity to synthesize chitin oligomers by some HASs, like the frog *Xenopus laevis* HAS^49,51^, SeHAS^26^ and *Mus musculus* HAS1^52^, supports the evolutionary connection between the synthesis of chitin and HA.

The phylogenetic reconstruction of fungal HASs, together with other Kingdom HASs, as well as the CHSs encoded by the same species that encode putative HASs and cellulose synthases (CSs) from *Arabidopsis thaliana*, *Komagataeibacter xylinus*, and *Novacetimonas hansenii* indicated a closer relationship between HASs and class VI CHSs (Figure 8C). HASs and CHSVI shared the same ancestor with class I, II and III CHSs. All these shared the same ancestor with analyzed CSs (Figure 8C). These results indicate that HASs are a direct innovation from a subset of CHSs; however, they are distant to CSs despite belonging to the same enzyme family, GT-2 processive glycosiltransferases^30^. Clustering analysis of HASs, CHSs and CSs showed a clear separation between CSs and the HAS-CHS cluster (Figure 8D), which confirmed that HASs are evolutionarily closer to CHSs than CSs.

## Discussion

### Yes, there are hyaluronic acid synthases in the Kingdom Fungi

*C. neoformans* CPS1p (CnHAS) is the only *bona fide* HAS reported so far^8^. However, in this work, 68 new putative HASs in 64 fungal species were found providing evidence that HASs might be more abundant in the Kingdom Fungi than expected (Figure 2).

Besides the conservation of catalytic motifs characteristic of GT2 processive glycosiltransferases (QXXRW, DXD and WGTR), they also strictly conserved the motifs essential for activity in other Kingdoms HASs: SG, which act as substrate binding site scaffold; GDDX, part of the catalytic site; and KR, which influences the processivity (Agarwal 2019) (Figure S1 and S2, Table S2). These motifs were also conformationally conserved among different Kingdoms HASs, including Kingdom Fungi (Figure 3), which supports the conservation of their catalytic function among the latter.

As other class I HASs^18,48^, fungal HASs harbor a single substrate binding site. The interaction of previously mentioned conserved motifs in CnHAS, and the other fungal HASs, with substrates UDP-GlcNAc and UDP-GlcUA was confirmed by docking analysis (Figure 4). R281 and W282 from the QXXRW motif, the three amino acids from the DXD motif, and W407 and R410 from the WGTR motif, interacted with the UDP-substrates, while R410 interacted only with UDP-GlcNAc, as previously reported^18,48^. D241 of GDDX motif also interacted with both substrates, while K242 from the same motif that harbors the catalytic function, interacted with UDP-GlcNAc^48^. Moreover, S205 from the SG motif and the R207, close to the latter, also interacted with both substrates^18^. Only the CnHAS KR motif did not interact with the substrates under the docking parameters, but they were in close proximity to them, thus a function for this motif cannot be ruled out. Additionally, L184 and R187 of the not fully conserved RYxxxFxxxR motif^18^ interacted with both substrates (Figure 4).

Besides conservation of a single catalytic site for both substrates within CnHAS, the 64 fungal species that encoded *has* genes also contained the machinery for the synthesis of HA precursors (Figure 2), which further supports the catalytic identity of *has* genes. HAS C (UDP-glucose pyrophosphorylase) catalyzes a central metabolism reaction, while HAS D (UDP N-acetyl glucosamine pyrophosphorylase) catalyzes the synthesis of UDP N-acetylglucosamine, precursor of chitin, a critical component of the fungal cell wall^53^; thus, ubiquitous presence of HAS C and HAS D in fungi is not surprising (Figure 1). However, HAS B catalyzes the synthesis of UDP-GlcUA, which is required only by fungi that synthesize polyuronides such as HA. In contrast, several putative HASs among Ascomycota, predicted by the AI CLEAN (Figure 2), did not encode for HAS B, one of the reasons why they were discarded for further analysis.

Artificial intelligence CLEAN platform was also utilized to further support fungal HASs identification. It managed to detect most of the HASs identified with HMM and strict conservation of critical motifs among Basidiomycetes, plus additional HASs among Ascomycota (Figure 2). Nevertheless, CLEAN-detected HASs systematically output low F1 scores (F1<0.6), which indicated low prediction reliability. Low F1 scores can be explained as CLEAN was trained with already characterized HASs, which led to low scores for the putative fungal ones. Despite CLEAN predictions, especially among species in the phylum Ascomycota, we parsimoniously preserved only the 68 sequences that strictly conserved the critical motifs of previously characterized HASs for further analysis, mainly found among Basidiomycetes. Those identified in Ascomycetes did not strictly conserve critical motifs, lacked the intrinsically disordered C-terminus, and the species that encoded them not always encoded HAS B. Thus, they were excluded, but they will be reserved for further analysis in the future.

### Putative fungal HASs were mainly found among Basidiomycetes

Fungal HASs were identified in Chytridiomycota, Mucoromycota, Ascomycota and Basidiomycota; nevertheless, it was in the latter where they were more abundant (Figure 2). Even within Basidiomycota, HASs were not evenly distributed. There, they appeared mainly in Tremellales, Boletales, Agaricales and Polyporales species (Figure 2). Basidiomycota species that encoded putative HASs inhabit a variety of heterogeneous ecological niches (saprophytes, mycorrhizas, vertebrate’s and plant’s pathogens, and endophytes) that make difficult to associate encoding of *has* genes with a particular environmental necessity. Also, those species have a diverse morphology and developmental patterns, some are yeast, others filamentous fungi and some others develop fruiting bodies. Therefore, the ecological niche, morphology and development cannot explain the abundance of putative HASs in Basidiomycota. Considering the different role of HA in bacteria and animals^54–56^, most probably HASs encoded by plant and animal pathogenic fungi form pericellular capsules, as *C. neoformans* HAS do^8^, useful in infective process that requires adhesion and immunological evasion^57^. Nevertheless, a role in development similar to animal HA cannot be ruled out among *has* gene encoding fungi that develop fruiting bodies.

### Despite necessary coincidences, fungal HASs are divergent: the transmembranal pore of putative fungal HASs was formed by only three alpha-helices

Despite the transmembranal pore of CnHAS is formed by three alpha-helices, instead of 4 that had been reported for bacterial HASs^18^ and 6 for vertebrates^48^ and viral HASs^20^, the docking analysis indicated that the pore had the biochemical assets necessary for translocation of HA. As previously reported for XlHAS pore^48^, the molecular docking analysis showed that CnHAS has a collar rich of positively charged amino acids at the base of the pore, followed by a ring of hydrophobic amino acids (Figure 5). However, it lacked the third ring of polar and hydrophobic amino acids present at the end of the pore of XlHAS. Although, CnHAS pore did not conserve the same amino acids identity than XlHAS pore, they conserved the same biochemical characteristics, which makes possible the translocation of HA through the pore conformed by only 3 helices. Furthermore, some amino acids that have been characterized in bacterial HASs pore also were identified in the docking model of CnHAS: residues K48 in TMH2 and E327 in TMH4 of SeHAS^39^, which influence the size of synthesized HA, were conserved in equivalent positions of CnHAS TMH2 and TMH4 such as R29 and D313, respectively; R205 and E213 of the polymer binding site RYxxxFxxxR of SeHAS^18^, was also present in the CnHAS model as R183 and E190, which interacted with the modeled HA hexamer. Despite convergent conserved features, the impact of observed differences in fungal HASs pore on the enzyme processivity needs to be empirically assessed.

### Fungal HASs gating loop is atypically connected to an intrinsically disordered region, a new regulatory mechanism?

The gating loop is a vital element for HASs activity across Kingdoms since it regulates the entrance of substrates to the catalytic site^48^. In bacterial HASs (SeHAS), truncation of the C-terminus, that includes the WGTR motif, leads to a significant decrease in enzymatic activity. Mutation of the residues of the gating loop motif, or in its proximity, affect the size of produced HA^58^. In XlHAS and CvHAS, the same phenomenon was observed when key amino acids from the WGTSGRK/R and WGTR motifs, respectively, were mutated^48^. Therefore, similar roles could be attributed to the gating loop present in fungal HASs due to conservation of the motif characterized in SeHAS and CvHAS, as well as the structural similarity to the SeHAS gating loop (Figure 6). Additionally, in the molecular docking simulation, UDP-sugars interacted with residues of this structure: the UDP bound to GlcNAc interacted with W407 and R410, and the UDP bound to GlcUA interacted with W407; in addition, the UDP-sugars interacted with R281 and W282 of the QXXRW motif (Figure 4), as previously reported for XlHAS^48^.

Even though the gating loop is conserved in all HASs, in bacterial HASs it is located at the very C-terminus, while in animal and viral HASs it is connected to an alpha helix of the transmembranal pore that provides stability to the gating loop conformation^48^. Among fungal HASs, the gating loop is connected to an intrinsically disordered region (IDR) whose size (between 30 and 200 amino acids) suggests a relevant biological function^59^. IDR are omnipresent in eukaryotic proteomes^60^. After a disorder-to-order transition, they are usually involved in protein-protein interactions that regulate a number of signaling pathways^61,62^. We hypothesize that the fungal HASs C-terminal IDR could mediate HASs interactions with different HASs protomers or regulatory proteins that could modulate the gating loop conformation and, ultimately, the HAS activity. This phenomenon has already been reported for a number of proteins such as the K+ channel KtrAB of *Vibrio alginolyticus* where an N-terminal IDR regulates the opening of the channel gating region^63^. Another case is exemplified by *Staphylococcus aureus* sortase A whose intrinsically disordered β6/β7 loop, after a disorder-to-order transition mediated by Ca^2+^, increases the enzyme activity^64^. Nevertheless, the hypothesis about the function of C-terminal IDR of fungal HASs should be experimentally tested.

Based on the connectivity of the gating loop and the composition of the transmembranal pore, we propose a new sub-classification for Class I HASs (Tabla 2): α-HAS, here are enclosed the vertebrates and viral HASs (CvHAS) whose gating loop is connected to IFH3 and TMH5; β-HAS, consisting of bacterial HASs whose gating loop is connected to IFH3 and the C-terminal end; and γ-HAS, where fungal HASs are enclosed, whose gating loop is connected to IFH3 and an IDR. The oligomerization was not the principal criterion for the subclassification because its inconsistency among vertebrates HASs: human and mouse HASs are present in an oligomeric form^23,24^; in contrast, XlHAS is in monomeric form^48^.

### What is the origin of fungal HASs?

It has been proposed that HASs have appeared independently several times in evolution^27,50^. The different directionality of HA synthesis among HASs supports this hypothesis. While vertebrates HASs elongate the HA at the reducing end^65,67^, bacterial HASs do it at the non-reducing end^21,68^, which implies different catalytic mechanisms to synthesize the same product using the same fold. Our own results are an expression of such diversity. In addition to some different conserved motifs among fungal HASs (Figure S2), the transmembranal pore is also differently constructed: while vertebratés HASs pore is composed of 6 helices, bacterial HASs one is built by 4 helices, and fungal HASs bear a minimalistic pore composed by only 3 helices (Figure 5). Moreover, the gating loop configuration highlights potential regulatory differences in a region conformationally sensitive and critical for enzyme catalysis and processivity^58^.

Despite chondroitin and chitin synthases have been proposed as the origin of HASs, the plasma membrane synthesis of HA, as well as the capacity of HASs to synthesize chitin oligomers strengthen the hypothesis of a CHS origin for HASs^50^. Lee and Spicer (2000)^33^ speculated that a B3 transferase activity was added to a previously existing B4 transferase activity from a CHS or CS, resulting in the creation of HASs. DeAngelis (2002)^17^ suggested that a CHS isoenzyme in an organism closely preceding chordates was mutated to create animal HASs. In Kingdom Fungi, since HASs are present in both early divergent fungi (Chrytidiomicota and Mucuromycota) and dykaria (Basidomycota and Ascomycota), HAS activity seems to have appeared early in fungi and as a product of independent convergent evolution, as suggested by the particularities of fungal HASs.

Our results suggest that fungal HASs share an ancestor with class VI CHS, and are close to class I, II and III CHSs (Figure 8D). Interestingly, class VI CHSs bear a single CON1 domain (PF03142), and have the simplest domain structure among CHSs^69^. Moreover, class VI CHSs are the only members of Division 3 CHSs and they are presumed the products of an independent CHS ancestor^70^. Interestingly, class VI CHSs are encoded by Ascomycetes (Pezizomycotina), several Chytridiomycota species, but they are almost completely absent among Mucoromycota and Basidiomycota species^70,71^, which, in contrast, encode HASs (Figure 6). Although further analyses are necessary, we speculate that, given the similarity and evolutionary closeness between fungal HASs and class VI CHSs, the common ancestor of both diverged to class VI CHSs in some fungi (Ascomycota), while in others it evolved to HASs (early divergent fungi and Basidiomycota).

## Materials and methods

### Reference HASs retrieval and analysis

In order to find HASs reference sequences (E.C. 2.4.1.212), as well as orthologs of enzymes that catalyze the synthesis of HA precursors: UDP-glucose-6-dehydrogenase (HAS B, EC 1.1.1.22), UDP-glucose pyrophosphorylase (HAS C, EC 2.7.7.9), and UDP-N-acetylglucosamine pyrophosphorylase (HAS D, EC 2.7.7.23) in Kingdom Fungi, a bibliographic search of already characterized HAS, HAS B, HAS C and HAS D was performed up to August 2023 (Table S1). Amino acid sequences of HAS, HAS B, HAS C and HAS D were retrieved either from UNIPROT^72^ or NCBI^73^.

HASs sequences were aligned in MUSCLE^74^ with default parameters and the alignments visualized and edited in JALVIEW^75^. Highly conserved amino acid residues along all the HASs sequences were identified and a database with those amino acids and their numbered position related to *C. neoformans* HAS (CPS1p) sequence was built (Table S2). HAS B, HAS C and HAS D orthologs were directly used to retrieve additional HAS B-D orthologs (see below).

### Search for putative fungal HAS, HAS B, HAS C and HAS D

To identify fungal HASs orthologs, *C. neoformans* CPS1p (CnHAS) sequence was used as a query to perform a locally run Blastp^76^, E=1x10^-20^, on a database that encompassed 910 fungal species proteomes obtained from NCBI^73^. Retrieved sequences were aligned with MUSCLE^74^ and edited in JALVIEW^75^. Only sequences that harbored all the essential residues and motifs of classical HASs (KR, DXD, R183, [S/T]G, T267, QXXR, WG[T/S]R) were selected for further analysis. From this data set (141 sequences), a Hidden Markov Model (HMM) profile was constructed in hmmbuild^77^ with default parameters and used to search again the fungal proteomes database with hmmsearch, E = 1x10^-20^. Recovered sequences (675) were aligned with MUSCLE and edited with Jalview. Sequences that did not contain characteristic HASs signatures were eliminated.

On the other hand, to retrieve additional fungal orthologs of HAS B, HAS C and HAS D, a multiple sequence alignment of previously identified and characterized reference sequences was made with MUSCLE^78^. Then, alignments were used to build HMM profiles with HMMER^79^, and used to search HAS B-D orthologs against our 910 fungal species proteomes database obtained from NCBI with hmmsearch, E-value of 1x10^-20^. Recursive searches were made until no more sequences were retrieved. Based on the data obtained from CLEAN and phylogenetic analysis (see below), a second filter was executed for HAS D (UDP-NAP), where only sequences harboring the motif LXXGGQGTRLGXXXPK^46,47^ were conserved.

Hmmsearch results for HAS, HAS B, HAS C and HAS D predicted orthologs were additionally assessed in CLEAN, contrastive learning–enabled enzyme annotation^40^, using default settings. Sequences predicted to be hyaluronic acid synthases (EC 2.4.1.212) were selected. For HAS B to D, F1 >0.6 was established as a cutoff parameter. Results of both HMM+conservation of typical HASs motifs, and CLEAN were grafted into a fungal tree of life reconstructed using the 910 reference fungi database with PhyloT, based on the NCBI taxonomy^80^. The tree was displayed with iTOL^81^.

### Phylogeny reconstruction

Phylogenies of retrieved orthologs were reconstructed with RaxML^82^ using LG as the best substitution model for all cases (HAS B, HAS C, HAS D, and fungal HAS) estimated using modelTest-ng allocated in RaxML^83^. HAS B, HAS C and HAS D orthologs from *S. equi subsp. Zooepidemicus* were used as outgroups (HAS B, ID: AEJ24424.1; HAS C, ID: AEJ24442.1, and HAS D, ID: AAQ05206.1). For fungal HAS, in addition to *S. equi subsp. Zooepidemicus*, *C. virus* HAS, and *H. sapiens* HAS2 were used as outgroup. Bootstrapping values were obtained after 350, 400 and 1000 iterations, respectively.

### Fungal HASs, CHSs and CS phylogenetic relationship

*In silico* identified CHSs^70^ from species encoding putative fungal HASs, as well as CSs from *Arabidopsis thaliana, Komagataeibacter xylinus,* and *Novacetimonas hansenii,* were retrieved from UNIPROT^72^. Reference HASs, putative fungal HASs, CHSs and *A. thaliana,* and *bacterial* CS were aligned with MUSCLE^74^ and manually edited in Jalview^75^. The alignment was edited to conserve only the central GT2 domain, deleting the N and C termini from position 69 to 282 using SeHAS sequence as reference. Phylogeny reconstruction of HASs, CHSs and CSs was performed in RaxML. The best-fitted substitution model (VT) was estimated with ModelTest-NG^83^ hosted in RaxML. Bootstrapping value was automatically estimated (auto-MRE).

### Clustering of HASs, CHSs and CSs

Clustering of HASs, CHSs and CSs was performed with EFI - Enzyme Similarity Tool^84^. Different alignment score thresholds (AST) were selected in order to visualize the clusters of interest: AST 144 to clusterize the putative and reference HASs; AST 6 to clusterize HASs, CHSs and CSs; AST 66 to clusterize HASs and CHSs. Results were analyzed and edited with Cytoscape^85^.

### Inference of secondary, and tertiary structures

Secondary structure of reference HASs (*C. neoformans* CPS1p, CnHAS; *S. equi* HAS, SeHAS; *X. laevis* HAS, XlHAS; and *H. sapiens* HAS2, HsHAS2) were predicted and visualized with Ali2D^86^ and 2dSS secondary structure visualization software^87^. TMHMM 2.0^88^ was used to infer transmembranal helices. Three-dimensional structures *of* CnHAS, SeHAS, XlHAS, HsHAS2, CcHAS and McHAS were predicted with AlphaFold^89^. HASs structural alignments were performed in Chimera 1.18^90^.

### Protein intrinsic disorder prediction

Intrinsically disordered regions of reference and fungal putative HASs were predicted through three different softwares: IUPRED3^91^, ESpritz 1.3^92^ and flDPnn^93^.

### Molecular docking studies

Docking simulations and subsequent analyses were performed in Chimera 1.18 using Autodock Vina^90,94^. 3D structures of wild type and mutant CnHAS were modeled in AlphaFold^89^. HA hexasaccharide was obtained from GLYCAM (glycam.org), while the UDP sugars were obtained from PubChem (https://pubchem.ncbi.nlm.nih.gov/). During simulations, the protein was kept rigid and the ligand flexible. In the analysis of the pore, the docking area was defined by a box of 35 x 55 x 45 Å, with the center in the intersection of axes perpendicular to one amino acid of each transmembranal helix (R22, L321 and K363). In the models with the UDP-sugars, the docking area was defined by a box of 35 x 40 x 25 Å with the center in K115, within the motif KR of the substrate binding site. The analysis was performed with the default settings, except that water was included. The energetically most favorable conformer was compared with the experimental model of XlHAS (PDB ID, 8smn)^48^. Docking analysis using relevant *in silico* mutants for the CnHAS pore and substrate binding site were run as negative controls.

## Acknowledgements

We thank the lab management work of Citlalli Rosales, IT assistance of Esteban Moreno, and the administrative support of Álvaro Zárate and CIATEJ staff.

This research was funded by Secretaría de Ciencia, Humanidades, Tecnología e Innovación (SECIHTI-Mexico), Ciencia de Frontera 2019-552259. LMF-H (CVU 1096158), MA-B (CVU 1167051) and PM-S (CVU 730994) were recipients of MSc or PhD fellowships from SECIHTI-Mexico.

## Conflict of interest

The authors declare that the research was conducted in the absence of any commercial or financial relationships that could be construed as a potential conflict of interest.

## Authors contribution

LMF-H curated already characterized HASs and identified new putative fungal HASs, performed tertiary and secondary structure prediction, estimated intrinsic disorder, and reconstructed initial phylogenies; MA-B curated already characterized HAS B-D and identified new putative fungal HAS B-D, complemented tertiary structural analysis and performed docking analysis; PM-S refined phylogenies and performed clustering analysis; EPP-M performed initial fungal HASs identification. LMF-H, MA-B, PM-S and JV designed experiments, analyzed results and wrote the manuscript. JV conceptualized research and acquired financial support. All authors critically reviewed and commented on the manuscript, and approved the final version.

## Abbreviations

HA: hyaluronic acid
HAS: hyaluronic acid synthase
CHS: chitin synthase
CS: cellulose synthase
IDR: intrinsically disordered region
UDP-GlcNAc: UDP-N-acetyl-D-glucosamine
UDP-GlcUA: UDP-glucuronic acid
HAS B, UDP-GDH: UDP-glucose 6-dehydrogenase
HAS C, UDP-GP: UDP-glucose pyrophosphorylase
HAS D, UDP-NAP: UDP-N-acetylglucosamine pyrophosphorylase
HMM: Hidden Markov Models
CnHAS: *Cryptococcus neoformans* HAS
SeHAS: *Streptococcus equi subsp. Zooepidemicus* HAS
HsHAS2: *Homo sapiens* HAS2
CvHAS: *Chlorella virus* HAS
CcHAS: *Coprinopsis cinerea* HAS
McHAS: *Mucor circinelloides* HAS
XlHAS: *Xenopus laevis* HAS

## Notes

### Competing Interest Statement

The authors have declared no competing interest.

