## Supporting Information for "Are hyaluronic acid synthases widely encoded in fungi?"

RUNNING TITLE: Fungal hyaluronic acid synthases

Laura Marina Franco-Herrera#, Mariandrea Aranda-Barba#, Paul Montaño-Silva, Eréndira Pérez-Muñoz and Jorge Verdín*

#These authors contributed equally to this work.

Biotecnología Industrial, CIATEJ-Centro de Investigación y Asistencia en Tecnología y Diseño del Estado de Jalisco, Zapopan, JAL, Mexico.

***Correspondence:**

Jorge Verdín, Biotecnología Industrial, CIATEJ. Camino Arenero 1227, Zapopan, JAL, México 45019. +52 (33) 3345-5200 ext. 2103

***Supportive figures***

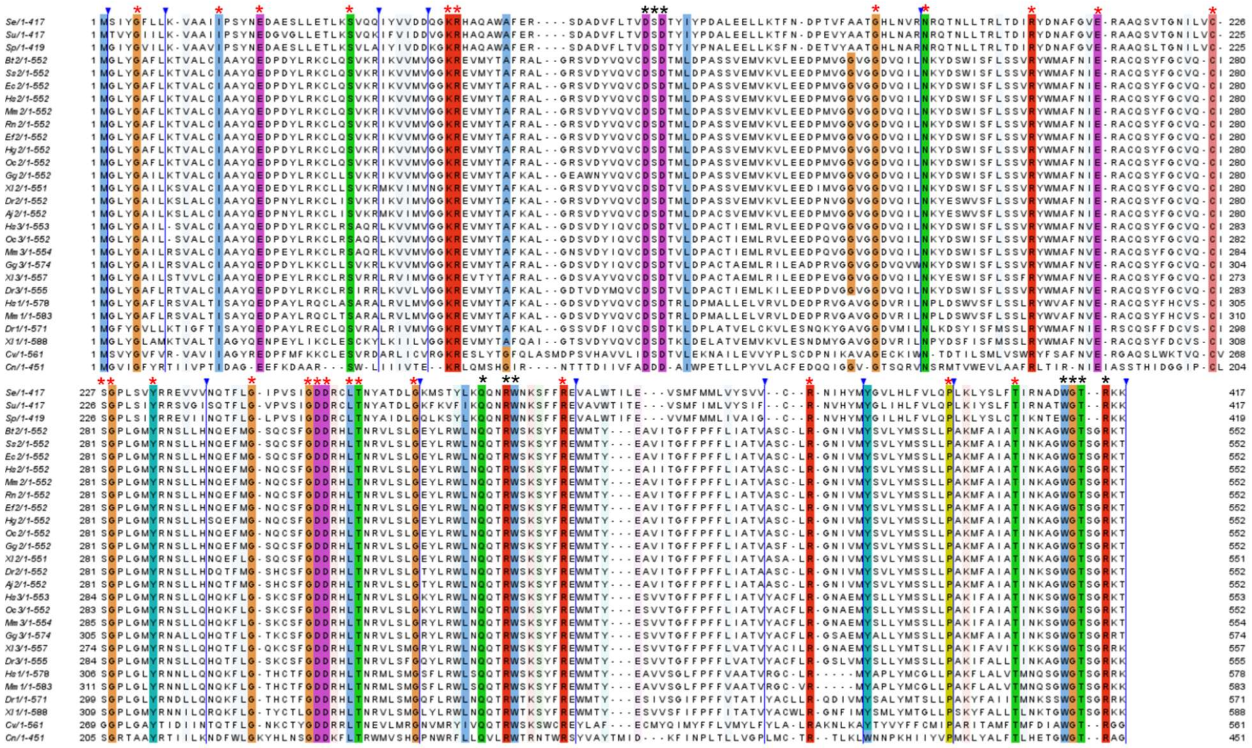

**Figure S1. Catalytically relevant amino acids are conserved among different Kingdoms HASs.** Amino acids sequence alignment of viral, bacteria, fungal and animal HASs. Characteristic amino acids of GT-2 glycosyltransferases are indicated with a black *, while amino acids that define HASs are indicated with a red *. Blue arrowheads and lines indicate sequence discontinuities where irrelevant amino acids were eliminated. Se, *Streptococcus equi* HAS; Su, *Streptococcus uberis* HAS; Sp, *Streptococcus pyogenes* HAS; Bt2, *Bos taurus* HAS2; Ss2, *Sus scrofa* HAS2; Ec2, *Equus caballus* HAS2; Hs2, *Homo sapiens* HAS2; Mm2, *Mus musculus* HAS2; Rn2, *Rattus norvegicus* HAS2; Ef2, *Eospalax fontanierii* HAS2; Hg2, *Heterocephalus glaber* HAS2; Oc2, *Oryctolagus cuniculus* HAS2; Gg2, *Gallus gallus* HAS2; Xl2, *Xenopus laevis* HAS2; Dr2, *Danio rerio* HAS2; Aj2, *Anguilla japonica* HAS2; Hs3, *H. sapiens* HAS3; Oc3, *O. cuniculus* HAS3; Mm3, *M. musculus* HAS3; Gg3, *G. gallus* HAS3; Xl3, *X. laevis* HAS3; Dr3, *D. rerio* HAS3;

Hs1, *H. sapiens* HAS1; Mm1, *M. musculus* HAS1; Dr1, *D. rerio* HAS1; Xl1, *X. laevis* HAS1; Cv, *Chlorella* virus HAS; Cn, *C. neoformans* CPS1p. The alignment was generated with Muscle[1] and edited with Jalview[2]. The figure was assembled in Inkscape.

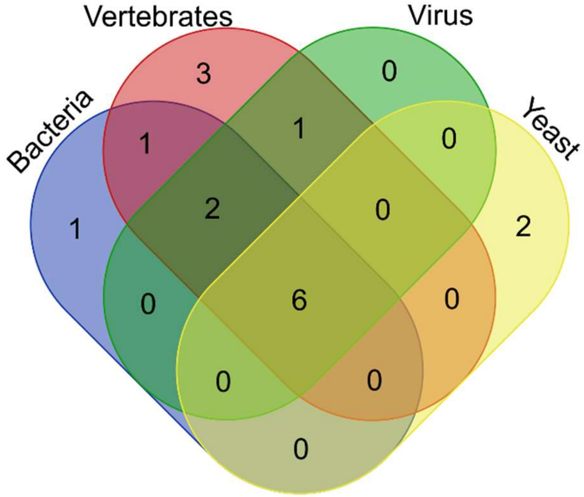


| Organism | Motifs |
|---|---|
| Bacteria / Vertebrates / Yeast / Virus | GDD KR QXXRW SG DXD WGTR |
| Bacteria / Vertebrates / Virus | KS RY |
| Bacteria / Vertebrates | AF |
| Vertebrates / Virus | TXL |
| Bacteria | TXI |
| Vertebrates | FA GXVGG WL |
| Yeast | GGVG RL |

**Figure S2. Shared and species specific conserved motifs of different Kingdoms HASs.** Despite all HASs shared characteristic motifs, some of them also harbored Kingdom specific signatures. Left, Venn diagram of the number of specific and shared motifs between bacteria, fungi, and animal (vertebrates) Kingdoms, and viral HASs. Right, shared and Kingdom specific motifs in bacteria, fungi, animal (vertebrates) and viral HASs.

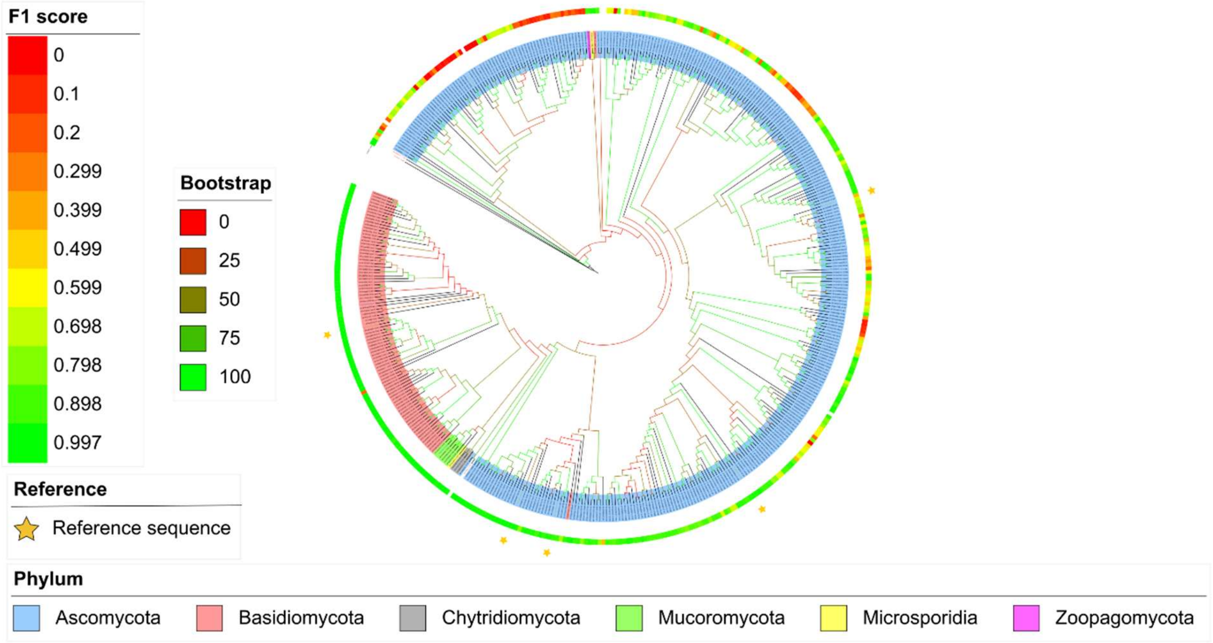


**Figure S3. Phylogeny reconstruction of HAS B fungal orthologs.** Departing from five already characterized UDP-glucose-6-dehydrogenase (HAS B, outermost yellow stars*)*, the phylogeny of HMM retrieved HAS B fungal orthologs was reconstructed in RaxML (best substitution model, LG); bootstrap was obtained after 350 cycles. For CLEAN predictions, F1 score heatmap is shown (external circle). Tree branches are colored after bootstrapping values.

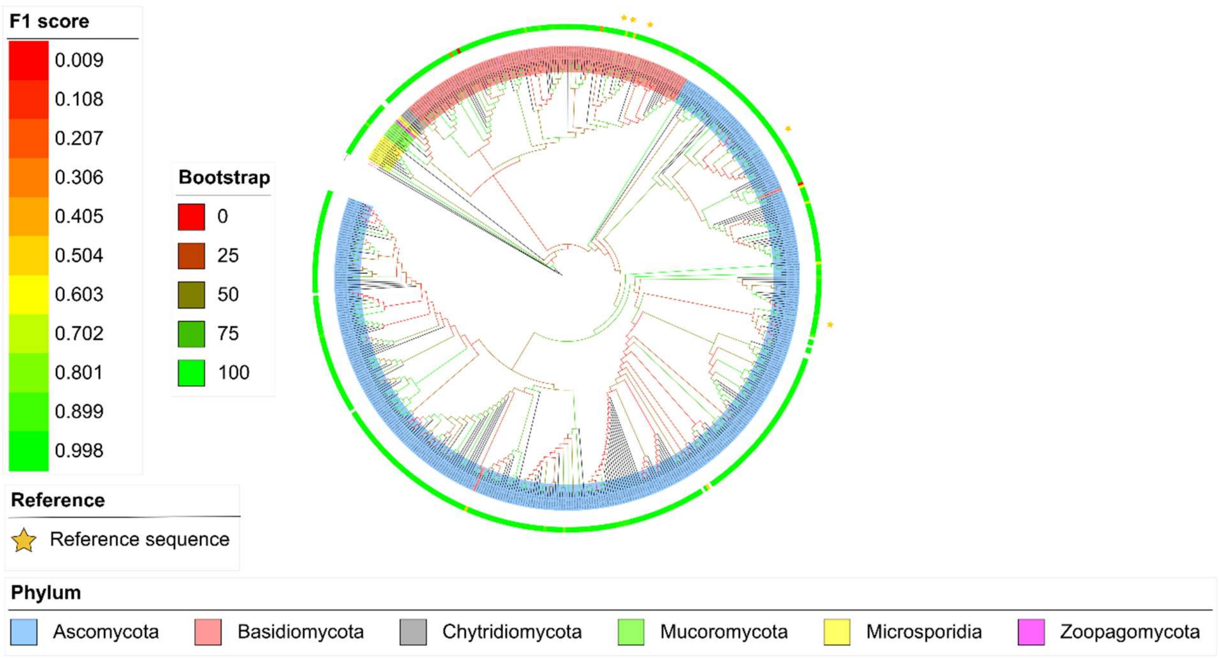


**Figure S4. Phylogeny reconstruction of HAS C fungal orthologs.** Departing from five already characterized UDP-glucose pyrophosphorylase (HAS C, outermost yellow stars*)*, the phylogeny of HMM retrieved HAS C fungal orthologs was reconstructed in RaxML (best substitution model, LG); bootstrap was obtained after 400 cycles. For CLEAN predictions, F1 score heatmap is shown (external circle). Tree branches are colored after bootstrapping values.

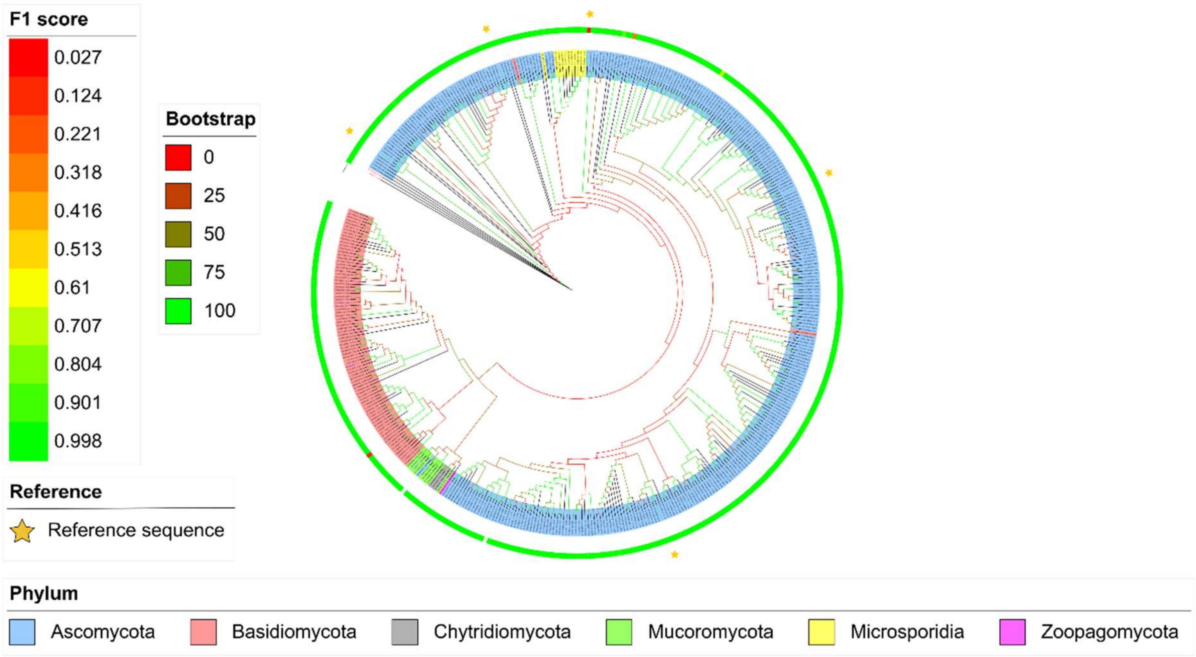


**Figure S5. Phylogeny reconstruction of HAS D fungal orthologs.** Departing from five already characterized UDP N-acetylglucosamine pyrophosphorylase (HAS D, outermost yellow stars*)*, the phylogeny of HMM retrieved HAS D fungal orthologs was reconstructed in RaxML (best substitution model, LG); bootstrap was obtained after 350 cycles. For CLEAN predictions, F1 score heatmap is shown (external circle). Tree branches are colored after bootstrapping values.

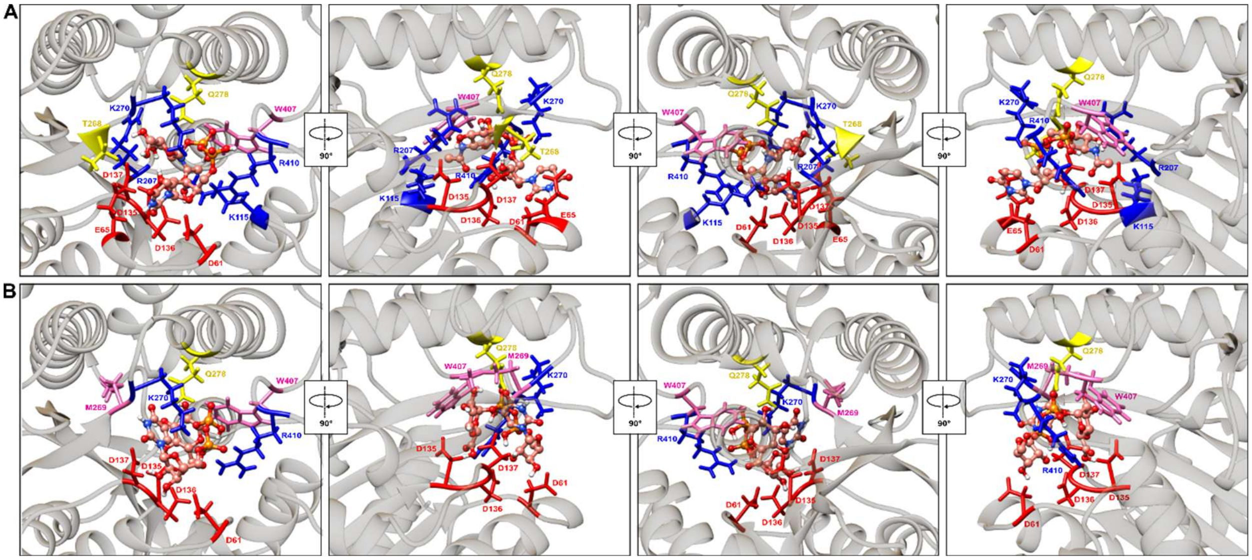


**Figure S6. Molecular docking analysis of modeled R281A/W282A *C. neoformans* HAS mutant and its substrates, UDP-GlcNAc and UDP-GlcUA.** A) Orthogonal views (90 ° rotation on "Y" axis) of the molecular docking analysis between modeled R281A/W282A *C. neoformans* HAS mutant and UDP-GlcNAc as substrate. The UDP-sugar complex interacted with D61, E65, K115, D135, D136, D137, R207, T268, K270, Q278, W407 and R410. B) Orthogonal views (90 ° rotation on "Y" axis) of molecular docking analysis between modeled R281A/W282A *C. neoformans* HAS mutant and UDP-GlcUA as substrate. The UDP-GlcUA interacted with D61, D135, D136, D137, M269, K270, Q278, W407 and R410. CnHAS R281A/W282A mutant led to a different interaction pattern with both substates compared to that of the wild type protein. CnHAS R281A/W282A mutant was modeled in AlphaFold 3[3]; docking simulations and subsequent analyses were performed in Chimera 1.18 using Autodock Vina[4].

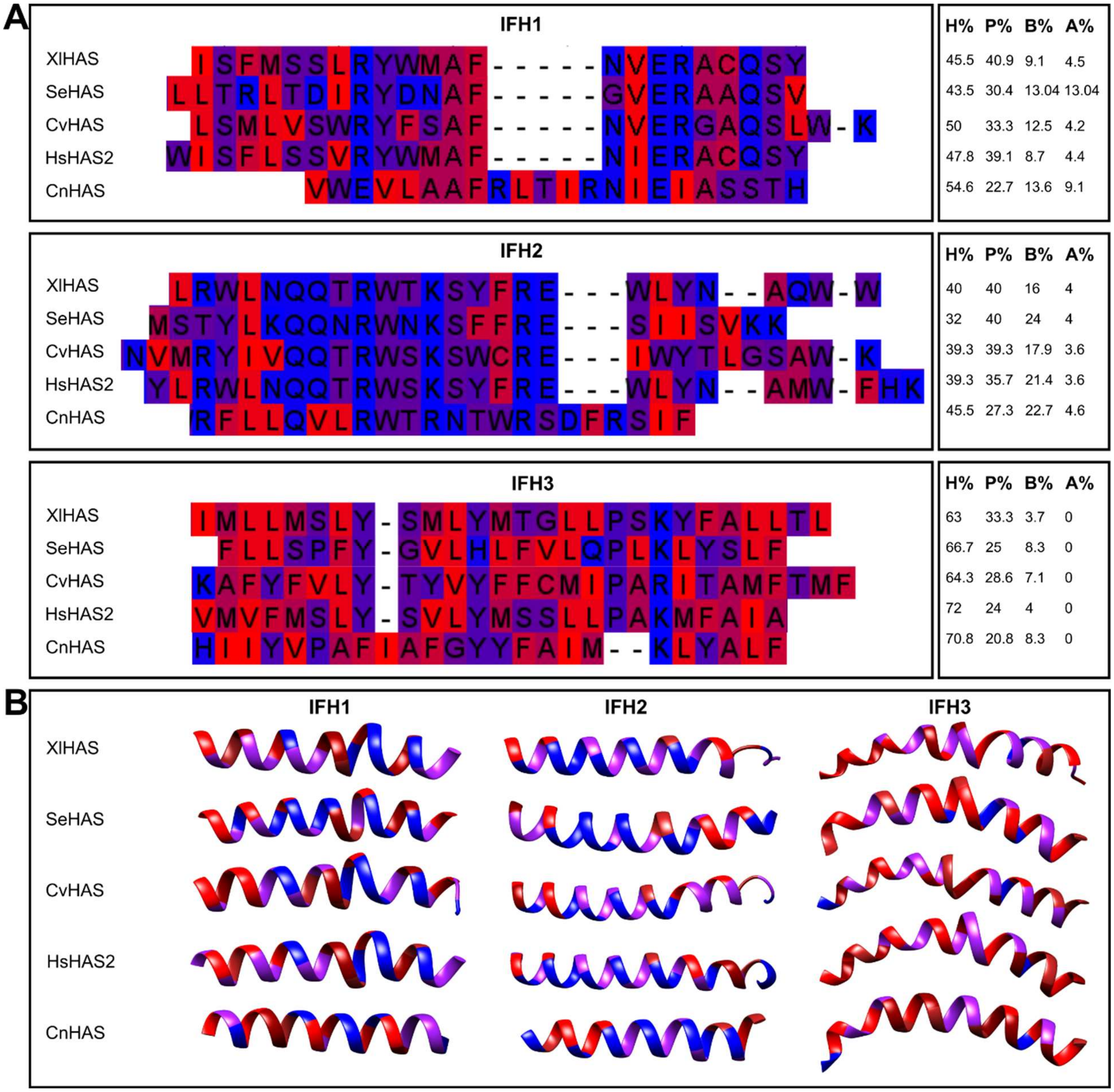


**Figure S7**. **Interface helices are strictly conserved among different Kingdoms HASs.** Besides their structural and spatial conservation, sequences of interface helices IFH1, IFH2 and IFH3 were conserved among viral, bacterial, fungal and animal HASs. A) In each interface helix (IFH1, IFH2 and IFH3), the hydrophobicity pattern is conserved (as darker red the square of the amino acid, more hydrophobic is) and also the percentage of each biochemical type of amino acid. H, hydrophobicity; P, polarity; B, basic, and A, acidic amino acids. B) Tridimensional structure of each IFH with the same color pattern as of hydrophobicity shown in A.

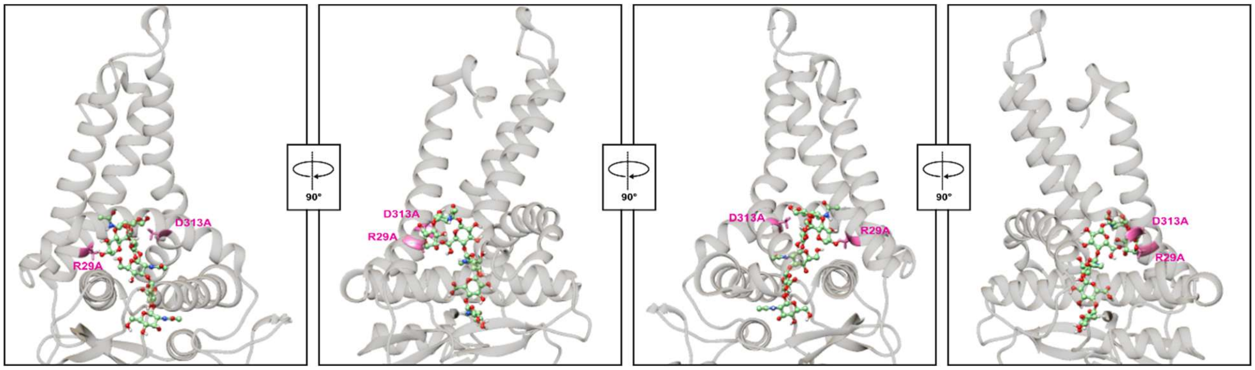


**Figure S8. Molecular docking analysis of modeled R29A/D313A *C. neoformans* HAS mutant and an HA oligomer.** Orthogonal views (90 ° rotation on "Y" axis) of the molecular docking analysis between modeled R29A/D313A *C. neoformans* HAS mutant and an HA oligomer. The HA oligomer interacted with the amino acids of the base of the pore (R22, W25, A29, D313A, K314, K366, F391 and K395) and some residues of the catalytic site (R183, I186, R187, E190, S205, R207, D241, K242, N285 and W282), but was unable to place itself within the pore. CnHAS R29A/D313A mutant was modeled in AlphaFold 3[3]; docking simulations and subsequent analyses were performed in Chimera 1.18 using Autodock Vina[4].

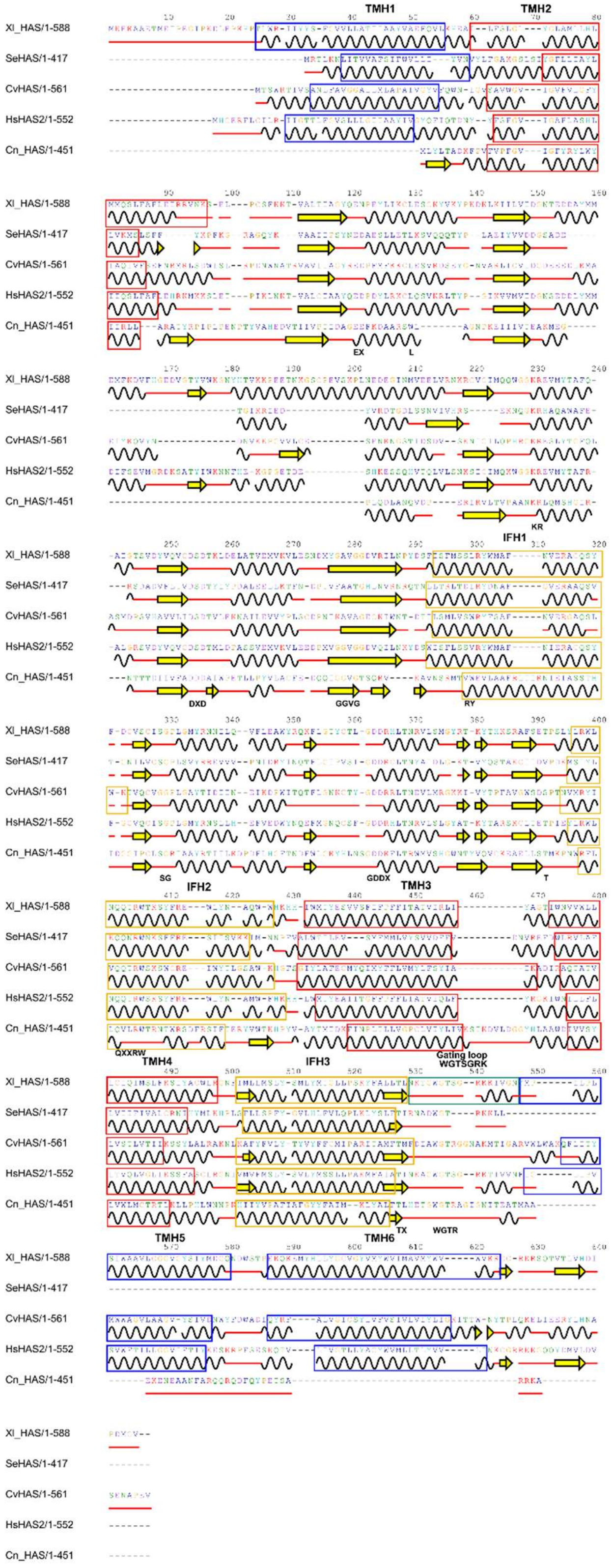
Xl_HAS/1-588
SeHAS/1-417
CvHAS/1-561
HsHAS2/1-552
Cn_HAS/1-451
TMH1
TMH2
EX
L
KR
IFH1
DXD
GGVG
RY
SG
GDDX
T
IFH2
TMH3
QXXRW
TMH4
IFH3
Gating loop
WGTSGRK
TMH5
TMH6
TX
WGTR

**Figure S9. Conservation and divergence of secondary structure elements among different Kingdoms HASs.** Secondary structure alignment of different Kingdoms HASs: *X. laevis*, *S. equi*, Chlorella virus, *H. sapiens* and *C. neoformans*. Although they conserved most secondary structure elements, bacterial HASs lacked helices H5 and H6, while fungal HASs were devoid of helices H1, H5 and H6. Alpha helices are represented as black waves, while beta-sheets are indicated as yellow arrows. Secondary structure prediction was performed in Ali2D[5] and visualized in 2dSS *secondary structure visualization*[6].

### *TABLES*

**Table S1.** Biochemically characterized HAS, HAS B, HAS C and HAS D used to identify putative fungal HASs.

**Table S2.** Function of conserved amino acids in *Streptococcus equi* and *Cryptococcus neoformans* HASs.

| Source | Se[1] numbering | Amino acid | Cn[2] numbering | Function | Reference |
|---|---|---|---|---|---|
| **Alignment** | **38** | **G** | **19** | - | - |
| Bibliography | 48 | K | 29 | Stability | [7] |
| Alignment | 67 | V | 55 | - | - |
| **Alignment** | **71** | **I** | **60** | - | - |
| Bibliography | 74 | Y | - | Interaction with precursor | [8] |
| **Bibliography** | **76** | **E** | **64** | **Interaction with precursor** | [8] |
| **Alignment** | **87** | **S** | **72** | - | - |
| Alignment | 98 | I | 81 | - | - |
| Alignment | 100 | V | 83 | - | - |
| Alignment | 101 | V | 84 | - | - |

| | | | | | |
|---|---|---|---|---|---|
| Bibliography | 103 | D | 95 | Initial binding with the substrate | [8] |
| **Bibliography** | **139** | **K** | **115** | **Essential for catalysis** | [8] |
| **Bibliography** | **140** | **R** | **116** | **Highly conserved** | [8] |
| **Bibliography** | **159** | **D** | **135** | **Stability** | [8] |
| **Bibliography** | **161** | **D** | **137** | **Stability** | [8] |
| Alignment | 164 | I | 139 | - | - |
| Alignment | 178 | D | 154 | - | - |
| **Alignment** | **186** | **G** | **161** | - | - |
| **Alignment** | **192** | **N** | **170** | - | - |
| Alignment | 201 | L | 179 | - | - |
| **Bibliography** | **205** | **R** | **183** | **Interaction with polysaccharide** | [8] |
| Bibliography | 206 | Y | 184 | Stability | [8] |
| Alignment | 212 | V | 189 | - | - |
| **Alignment** | **213** | **E** | **190** | - | - |
| Alignment | 217 | Q | 194 | - | - |
| Bibliography | 218 | S | 195 | - | [8] |
| Alignment | 223 | I | 201 | - | - |
| **Bibliography** | **226** | **C** | **203** | - | [8] |
| Bibliography | 227 | S | 205 | Stability | [8] |
| **Bibliography** | **228** | **G** | **206** | **Provides flexibility to substrate-interacting loop** | [8] |
| **Bibliography** | **233** | **Y** | **211** | **Influence on activity** | [8] |
| Bibliography | 234 | R | 212 | Influence on activity | [8] |
| **Alignment** | **250** | **F** | **230** | - | - |
| Alignment | 251 | L | 231 | - | - |
| **Alignment** | **252** | **G** | **232** | - | - |
| **Bibliography** | **258** | **G** | **239** | - | [8] |
| **Bibliography** | **259** | **D** | **240** | **Influence on activity, substrate interaction** | [8] |
| **Bibliography** | **260** | **D** | **241** | **Influence on activity** | [8] |
| Bibliography | 261 | R | 242 | Influence on activity | [8] |

| | | | | | |
|---|---|---|---|---|---|
| **Bibliography** | **263** | **L** | **244** | **Influence on activity** | [8] |
| **Alignment** | **264** | **T** | **245** | - | - |
| **Alignment** | **271** | **G** | **252** | - | - |
| Bibliography | 281 | C | 260 | Influence on activity | [8] |
| Bibliography | 283 | T | 268 | Essential for catalysis | [8] |
| Alignment | 292 | Y | 275 | - | - |
| Alignment | 293 | L | 276 | - | - |
| **Bibliography** | **295** | **Q** | **278** | **Essential for catalysis** | [8] |
| **Bibliography** | **298** | **R** | **281** | **Essential for catalysis** | [8] |
| **Bibliography** | **299** | **W** | **282** | **Essential for catalysis** | [8] |
| Alignment | 301 | K | 284 | - | - |
| Alignment | 302 | S | 285 | - | - |
| **Alignment** | **305** | **R** | **288** | - | - |
| Alignment | 323 | W | 309 | - | - |
| Bibliography | 327 | E | 313 | Important for processivity | [7] |
| Bibliography | 367 | C | 361 | Not essential | [8] |
| **Alignment** | **368** | **R** | **363** | - | - |
| Alignment | 386 | Y | 372 | - | - |
| **Alignment** | **396** | **P** | **382** | - | - |
| Alignment | 398 | K | 395 | - | - |
| **Alignment** | **404** | **T** | **401** | - | - |
| Alignment | 405 | I | 402 | - | - |
| **Bibliography** | **410** | **W** | **407** | **Essential for catalysis. Gating loop** | [9] |
| **Bibliography** | **411** | **G** | **408** | **Gating loop** | [9] |
| **Bibliography** | **412** | **T** | **409** | **Gating loop** | [9] |
| **Bibliography** | **413** | **R** | **410** | **Important for processivity. Gating loop** | [9] |

[1] *Streptococcus equi* HAS

[2] *Cryptococcus neoformans* HAS

| | |
|---|---|
| (green) | Conserved in >90% sequences |
| (yellow) | Function reported in literature, but not fully conserved |
| **BOLD RED** | Conserved in 100% sequences |

**TABLE S3.** Putative fungal hyaluronic acid synthases*

| PROTEIN ID | Organism | CLEAN functional prediction/F1 score |
|---|---|---|
| EPB81503.1 | *Mucor circinelloides 1006PhL* | EC:2.4.1.212/0.0208 |
| XP_001830918.2 | *Coprinopsis cinerea okayama7#130* | EC:2.4.1.212/0.0090 |
| XP_001877942.1 | *Laccaria bicolor S238N-H82* | EC:2.4.1.212/0.0056 |
| XP_003029223.1 | *Schizophyllum commune H4-8* | - |
| XP_003195746.1 | *Cryptococcus gattii WM276* | EC:2.4.1.212/0.0123 |
| XP_006457289.1 | *Agaricus bisporus var. bisporus H97* | - |
| XP_006676814.1 | *Batrachochytrium dendrobatidis JAM81* | - |
| XP_007001154.1 | *Tremella mesenterica DSM 1558* | - |
| XP_007266631.1 | *Fomitiporia mediterranea MF3/22* | EC:2.4.1.212/0.0220 |
| XP_007309723.1 | *Stereum hirsutum FP-91666 SS1* | EC:2.4.1.212/0.0135 |
| XP_007318053.1 | *Serpula lacrymans var. lacrymans S7.9* | EC:2.4.1.212/0.0241 |
| XP_007333788.1 | *Agaricus bisporus var. burnettii JB137-S8* | - |
| XP_007381009.1 | *Punctularia strigosozonata HHB-11173 SS5* | EC:2.4.1.212/0.0427 |
| XP_007404447.1 | *Melampsora larici-populina 98AG31* | - |
| XP_007767154.1 | *Coniophora puteana RWD-64-598 SS2* | - |
| XP_007864082.1 | *Gloeophyllum trabeum ATCC 11539* | EC:2.4.1.212/0.0206 |
| XP_008034657.1 | *Trametes versicolor FP-101664 SS1* | EC:2.4.1.212/0.0215 |
| XP_009546987.1 | *Heterobasidion irregulare TC 32-1* | EC:2.4.1.212/0.0227 |
| XP_012051731.1 | *Cryptococcus neoformans var. grubii H99* | EC:2.4.1.212/0.0248 |
| XP_012177469.1 | *Fibroporia radiculosa* | EC:2.4.1.212/0.0335 |
| XP_013242036.1 | *Tilletiaria anomala UBC 951* | - |
| XP_014179419.1 | *Trichosporon asahii var. asahii CBS 2479* | - |
| XP_014565991.1 | *Mixia osmundae IAM 14324* | - |
| XP_016611856.1 | *Spizellomyces punctatus DAOM BR117* | - |
| XP_018260836.1 | *Kwoniella dejecticola CBS 10117* | - |
| XP_018282379.1 | *Cutaneotrichosporon oleaginosum]* | - |
| XP_018293447.1 | *Phycomyces blakesleeanus NRRL 1555(-)* | - |
| XP_018294775.1 | *Phycomyces blakesleeanus NRRL 1555(-)* | - |
| XP_018989215.1 | *Cryptococcus amylolentus CBS 6039* | EC:2.4.1.212/0.0121 |
| XP_019001392.1 | *Kwoniella mangroviensis CBS 8507* | - |
| XP_019010367.1 | *Kwoniella pini CBS 10737* | - |
| XP_019031048.1 | *Cryptococcus wingfieldii CBS 7118* | EC:2.4.1.212/0.0336 |

| | | |
|---|---|---|
| XP_019044101.1 | *Kwoniella bestiolae CBS 10118* | - |
| XP_021874209.1 | *Kockovaella imperatae* | EC:2.4.1.212/0.0179 |
| XP_021876649.1 | *Lobosporangium transversale* | EC:2.4.1.212/0.0188 |
| XP_021881596.1 | *Lobosporangium transversale* | EC:2.4.1.212/0.0684 |
| XP_023465919.1 | *Rhizopus microsporus ATCC 52813* | EC:2.4.1.212/0.0041 |
| XP_024342638.1 | *Postia placenta MAD-698-R-SB12* | EC:2.4.1.212/0.0108 |
| XP_025358400.1 | *Meira miltonrushii* | - |
| XP_025370536.1 | *Ceraceosorus guamensis* | - |
| XP_025377642.1 | *Acaromyces ingoldii* | - |
| XP_025600626.1 | *Tilletiopsis washingtonensis* | EC:2.4.1.212/0.0115 |
| XP_026624494.1 | *Aspergillus welwitschiae* | EC:2.4.1.212/0.0033 |
| XP_027611330.1 | *Sparassis crispa* | EC:2.4.1.212/0.0226 |
| XP_028473368.1 | *Apiotrichum porosum* | EC:2.4.1.212/0.0109 |
| XP_031026858.1 | *Synchytrium microbalum* | - |
| XP_031026966.1 | *Synchytrium microbalum* | - |
| XP_031861725.1 | *Kwoniella shandongensis* | - |
| XP_036630232.1 | *Pleurotus ostreatus* | EC:2.4.1.212/0.0089 |
| XP_037223101.1 | *Mycena indigotica* | - |
| XP_040631151.1 | *Dacryopinax primogenitus* | EC:2.4.1.212/0.0027 |
| XP_040761921.1 | *Laetiporus sulphureus 93-53* | EC:2.4.1.212/0.0261 |
| XP_041165207.1 | *Suillus plorans* | EC:2.4.1.212/0.0252 |
| XP_041190470.1 | *Suillus subaureus* | - |
| XP_041205320.1 | *Suillus clintonianus* | EC:2.4.1.212/0.0126 |
| XP_041216565.1 | *Suillus fuscotomentosus* | - |
| XP_041227617.1 | *Suillus fuscotomentosus* | EC:2.4.1.212/0.0141 |
| XP_041249170.1 | *Suillus subalutaceus* | EC:2.4.1.212/0.0150 |
| XP_041293822.1 | *Suillus discolor* | EC:2.4.1.212/0.0179 |
| XP_041310044.1 | *Suillus bovinus* | - |
| XP_041545200.1 | *Aspergillus luchuensis* | EC:2.4.1.212/0.0012 |
| XP_043014641.1 | *Marasmius oreades* | - |
| XP_046083831.1 | *Lentinula edodes* | EC:2.4.1.212/0.0022 |
| XP_047752471.1 | *Psilocybe cubensis* | EC:2.4.1.212/0.0010 |
| XP_047780802.1 | *Rhodofomes roseus* | - |
| XP_047874229.1 | *Epithele typhae* | EC:2.4.1.212/0.0022 |
| XP_047897365.1 | *Neoantrodia serialis* | EC:2.4.1.212/0.0128 |
| XP_572855.1 | *Cryptococcus neoformans var. neoformans JEC21* | EC:2.4.1.212/0.0168 |
| XP_773764.1 | *Cryptococcus neoformans var. neoformans B-3501A* | EC:2.4.1.212/0.0168 |

*Putative fungal hyaluronic acid synthases identified with hidden Markov models and depurated by conservation of catalytically essential amino acids. Putative HASs identified by CLEAN AI are also indicated.

**Table S4.** Docking analysis interactions between wild and mutant CnHAS and GlcNAc and GlcUA

| | Interactions with wild CnHAS | | | |
|---|---|---|---|---|
| **Amino acid** | **GlcNAc** | **GlcUA** | **Motif** | **Function** |
| **D135** | H bond, ionic bond | Ionic bond | DXD | Substrate binding |
| **D136** | Ionic bond | Ionic bond | | |
| **D137** | Ionic bond | Ionic bond | | |
| **Q164** | | H bond | | |
| **L184** | Hydrophobic interaction | Hydrophobic interaction | RYxxxFxxxR | Polymer binding |
| **R187** | Ionic bond | Ionic bond | | |
| **S205** | H bond | H bond | SG | Binding site scaffold |
| **R207** | Ionic bond | Ionic bond | | Close to SG |
| **D241** | Ionic bond | Ionic bond | GDDK | Catalytic base |
| **K242** | Ionic bond | | | |
| **M269** | | Hydrophobic interaction | | |
| **R281** | Hydrophobic interaction, ionic bond | Hydrophobic interaction, ionic bond | QXXRW | Substrate and polymer binding |
| **W282** | H bond (only UDP), hydrophobic interaction | H bond (only UDP), hydrophobic interaction | | |
| **W407** | Hydrophobic interaction | Hydrophobic interaction | WGTR | Gating loop |
| **R410** | Ionic bond | | | |

| | Interactions with R281A/W282A mutant CnHAS | | | |
|---|---|---|---|---|
| **Amino acid** | **GlcNAc** | **GlcUA** | **Motif** | **Function** |
| **D61** | Ionic bond | Ionic bond | | |
| **E65** | Ionic bond | | | |
| **K115** | Ionic bond | | KR | Substrate binding, charge neutralization |

| | | | | |
|---|---|---|---|---|
| **D135** | H bond, ionic bond | Ionic bond | | |
| **D136** | H bond, ionic bond | Ionic bond | DXD | Substrate binding |
| **D137** | Ionic bond | Ionic bond | | |
| **R207** | H bond, ionic bond | | | Close to SG motif |
| **T268** | H bond | | | Substrate binding |
| **M269** | | Hydrophobic interaction | | |
| **K270** | Ionic bond | H bond, ionic bond | | |
| **Q278** | H bond | H bond | QXXRW | Substrate and polymer binding |
| **W407** | Hydrophobic interaction | Hydrophobic interaction | | |
| **R410** | H bond, ionic interaction | H bond (only UDP), ionic interaction | Motif WGTR | Gating loop |

*The squares in light orange outline the amino acids that interact with the nucleotide of the UDP-sugar; the yellow square indicates the amino acids that interact with the sugar moiety of the UDP-sugar; and the orange square encloses the amino acids that interact with both elements of the UDP-sugar substrate.
